# Rapid Directed Molecular Evolution of Fluorescent Proteins in Mammalian Cells

**DOI:** 10.1101/2021.08.02.454744

**Authors:** Siranush Babakhanova, Erica E. Jung, Kazuhiko Namikawa, Hanbin Zhang, Yangdong Wang, Oksana M. Subach, Dmitry A. Korzhenevskiy, Tatiana V. Rakitina, Xian Xiao, Wenjing Wang, Jing Shi, Mikhail Drobizhev, Demian Park, Lea Eisenhard, Hongyun Tang, Reinhard W. Köster, Fedor V. Subach, Edward S. Boyden, Kiryl D. Piatkevich

## Abstract

*In vivo* imaging of model organisms is heavily reliant on fluorescent proteins with high intracellular brightness. Here we describe a practical method for rapid optimization of fluorescent proteins via directed molecular evolution in cultured mammalian cells. Using this method, we were able to perform screening of large gene libraries containing up to 2·10^7^ independent random genes of fluorescent proteins expressed in HEK cells completing one iteration directed evolution in a course of ∼8 days. We employed this approach to develop a set of green and near-infrared fluorescent proteins with enhanced intracellular brightness. The developed near-infrared fluorescent proteins demonstrated high performance for fluorescent labeling of neurons in culture and *in vivo* in model organisms such as *C.elegans*, *Drosophila*, zebrafish, and mice. Spectral properties of the optimized near-infrared fluorescent proteins enabled crosstalk-free multicolor imaging in combination with common green and red fluorescent proteins, as well as dual-color near-infrared fluorescence imaging. The described method has a great potential to be adopted by protein engineers due to its simplicity and practicality. We also believe that the new enhanced fluorescent proteins will find wide application for *in vivo* multicolor imaging of small model organisms.

## Introduction

Fluorescent proteins (FPs) became indispensable tools for *in vivo* imaging of cellular and subcellular structures in model organisms^1^. Since the cloning of the first green FP from jellyfish *Aequorea victoria* in 1992^2^, the myriads of chromoproteins with various spectral and biochemical properties have been cloned from diverse natural sources such as corals^3^, fish^4^, plants^5^, soil bacteria^6^, and cyanobacteria^7^. However, all naturally occurring chromoproteins have to be modified, optimized, or even re-engineered in order to be utilized for fluorescence imaging *in vivo*^8–12^. Among all biochemical characteristics, the intracellular brightness is one of the most crucial properties for *in vivo* applications that protein engineers and developers choose to optimize before everything else^13,14^. FPs are usually optimized via directed molecular evolution by iteratively generating and screening large gene libraries in bacterial cells^13–15^. However, high molecular brightness, commonly screened for in bacterial cells, does not always correspond to high intracellular brightness when expressed in cultured mammalian cells^8,9,16^ or *in vivo*^16–18^. This phenomenon is particularly well documented for bacteriophytochrome-based FPs^16,19,20^, whose fluorescence relies on the incorporation of the chromophore biliverdin (BV) from the bulk^21^. Ideally, the development of functional proteins for *in vivo* imaging should be performed in an environment physiologically relevant to the final hosts to ensure proper protein folding, localization, and posttranslational modifications^22,23^. Yeasts, although eukaryotic cells, which provide convenience of large gene library expression like bacteria, may not serve as an ideal host system for FPs as it was shown that brightness in yeast cells does not necessarily retain in mammalian cells^24^. In this regard, vertebrate cell lines represent a promising expression host for directed molecular evolution of FPs. Indeed, several studies demonstrated possibility to evolve FPs in cultured chicken^25^ or mammalian cells^26–28^. However, the proposed methods did not find wide adaptation among protein engineers due to several limitations and drawbacks. First, all previously developed methods for directed molecular evolution in cultured cells from vertebrates involve establishing cell lines stably maintaining target genes, in a way where any given cell expresses ideally no more than one copy of a target gene^26–28^. However, both commonly used single gene copy delivery methods, such as electroporation and retroviral transduction, and establishing stable cells lines are time-consuming and laborious, complicated by apoptosis, low efficiency of stable gene integration, and long cell doubling time. Second, *in situ* diversification of target genes using a cytidine de aminase^25,26,29^ or viral replication^30^ has a low mutation rate of only 1-3 mutations per kilobase pair in comparison to 9-16 mutations per kilobase pair for regular error-prone PCR typically used for FP development. Higher mutation rate can be achieved with CRISPR/Cas9 editing technology^28^ or via *in vitro* mutagenesis^27^, however, it comes with a limited library size of 10^5^-10^6^ independent clones. Third, recovery of target genes after screening and sorting is typically done from the large pools (10^3^-10^7^) of collected cells with subsequent random picking of just a few (10-200) clones from the pool that significantly reduces the chance of finding the best variant^26–28^. As a result, the previously reported methods of FP evolution in mammalian cells remained as proof of principle studies and did not yield any practically useful FPs for *in vivo* imaging.

Here we described a simple and practical method that overcomes major limitations and drawbacks of currently available methods for directed molecular evolution of FPs in mammalian cells. The developed method allowed us to perform optimization of intracellular brightness in the course of 4 days per iteration by generation and screening of large (up to 2×10^7^) and highly diverse (10-15 mutations per kilobase pair) libraries of FPs in mammalian cells and extra 3 days for isolation and identification of individual variants. Moreover, our method allows rapid target gene recovery from very small pools of collected cells (10-100 cells) enabling genotyping and detailed phenotyping of rare cell populations common for large random libraries. We employed the described method to develop a set of green and near-infrared (NIR) FPs, named phiLOV3, TagRFP658, and miRFP2, with enhanced intracellular brightness. The improvements of intracellular brightness for phiLOV3 and miRFP2 did not correlate with molecular brightness when compared to respective parental proteins thus highlighting importance of protein optimization in the corresponding cellular context. Using rational design, we generated an enhanced version of miRFP2 with greatly increased intracellular brightness, called emiRFP2. We demonstrated applicability of the developed proteins for *in vivo* imaging of neurons in *C.elegans*, *Drosophila*, zebrafish, and intact mouse brain tissue. Combination of the developed FPs with other common green and red FPs enabled crosstalk-free multicolor imaging using standard optical configurations. Moreover, we performed dual-color NIR imaging of subcellular structures using combination of emiRFP2 with blue-shifted NIR FP mCardinal. The developed method has a great potential to be adopted by protein engineers for optimization of FPs in mammalian cells. We also believe that the new enhanced FPs will find wide application for *in vivo* imaging of small model organisms.

## Results

### Protein optimization in mammalian cells

The rapid protein optimization via directed molecular evolution is based on a simple and scalable method for expression of large gene libraries in mammalian cells in combination with high-throughput live cell screening techniques, *e.g.*, fluorescence activated cell sorting (FACS). The workflow includes six major hands-on steps: i) preparation of gene library; ii) transfection of mammalian cells with gene library in bulk; iii) screening and collection of individual cells; iv) cloning of the target genes from selected cells into expression vector; v) transfection of the cloned plasmids into mammalian cells; vi) imaging and selection of individual clones (**Figure 1a**). One iteration of directed molecular evolution can be carried out in about 8 days (**Figure 1b**). In case of the bulk selection, when pools of selected genes are subjected to the next round of evolution, mutagenesis and screening can be performed with a period of ∼3.5 days (**Figure 1b**). To validate the proposed approach, we decided to enhance intracellular brightness of the biochemically and spectrally distinct FPs. We chose a set of FPs that originated from the four evolutionary different classes of chromoproteins and those that are diverse in their later synthetic evolution conditions. Namely, we chose a green FP phiLOV2.1 engineered from the flavin-binding LOV (light, oxygen, or voltage sensing) domain^31^, a naturally occurring green FP UnaG cloned from freshwater eel *Anguilla japonica*^4^, an engineered far-red fluorescent GFP-like FP TagRFP657 (*ref.*^32^), and a near-infrared FP miRFP derived from PAS-GAF domains of the *Rp*BphP1 bacteriophytochrome^33^. By selecting already optimized FPs as templates our goal was to demonstrate a great potential of the proposed approach to further enhance desired characteristics in mammalian cells. First, human-codon optimized genes of the selected proteins were subjected to error-prone PCR and cloned into the mammalian expression vector containing an SV40 origin of replication. The SV40 origin of replication significantly increases the expression levels of transgene under transient transfection conditions due to episomal replication of the plasmid within a host cell that expresses the SV40 large T-antigen^34^, which is crucial under one plasmid per cell transfection conditions. The generated random libraries, containing around 10^6^-10^7^ independent clones, were transfected into HEK293FT cells using the modified calcium phosphate method, which was optimized for single gene copy per cell delivery^35^. After plasmid replication and protein expression for 48 h, we used FACS to collect ∼100 cells with the highest fluorescence intensity for each library. Next, the target genes, recovered from the pools of collected cells, were either subjected to the next round of directed evolution or directly cloned into the mammalian expression vector (**Figure 1a**). In the latter case, several hundreds of randomly picked clones were individually transfected into HEK cells to compare their brightness to the corresponding parental proteins. Overall, one iteration of directed evolution was carried out within ∼3.5 days and additional ∼4 days were required to clone, express, assess, and sequence individual mutants (**Figure 1b**). We carried out two rounds of directed evolution for each template followed by screening of individually picked clones expressed in HEK cells under fluorescence microscope (see **Supplementary Table 1** for the screening conditions and parameters). Assessment of the clones selected from the UnaG library did not identify variants with improved intracellular brightness, although sequencing of the brightest selected variants revealed amino acid substitutions in the structurally important regions (**Supplementary Figure 1, 2**). Therefore, we did not perform further characterization of the UnaG mutants. For phiLOV2.1 and TagRFP657, we identified multiple variants that outperformed corresponding parental proteins in terms of intracellular brightness and in case of miRFP we identified only one mutant with improved brightness (**Supplementary Figure 1**). We selected the brightest variant from each library for further evaluation. However, during imaging we found out that the brightest TagRFP657 variant had very limited photostability, we decided to choose the second brightest variant, which was confirmed to have higher than the parental protein photostability (**Supplementary Figure 1**). To confirm that the observed improvements were statistically significant, we repeated the measurements for each selected variant on several biological replicates in HEK cells. Indeed, all variants showed biologically and statistically significant improvements in intracellular brightness. The phiLOV2.1 variant showed 2.8-fold higher brightness over parental protein, the TagRFP657 and miRFP variants showed 30% and 27% over parental proteins, respectively (**Figure 1c-e**). Sequence analysis of the selected variants revealed two amino acid mutations in phiLOV2.1, six in TagRFP657, and nine in miRFP (**Supplementary Figure 3-5**). Correspondingly, we named the identified variants as phiLOV3, TagRFP658, and miRFP2, and used them for further detailed characterization. As a result, we were able to complete two iterations of directed molecular evolution in mammalian cells within eight days enhancing intracellular brightness of the selected FPs (**Figure 1**).

**Figure 1.**
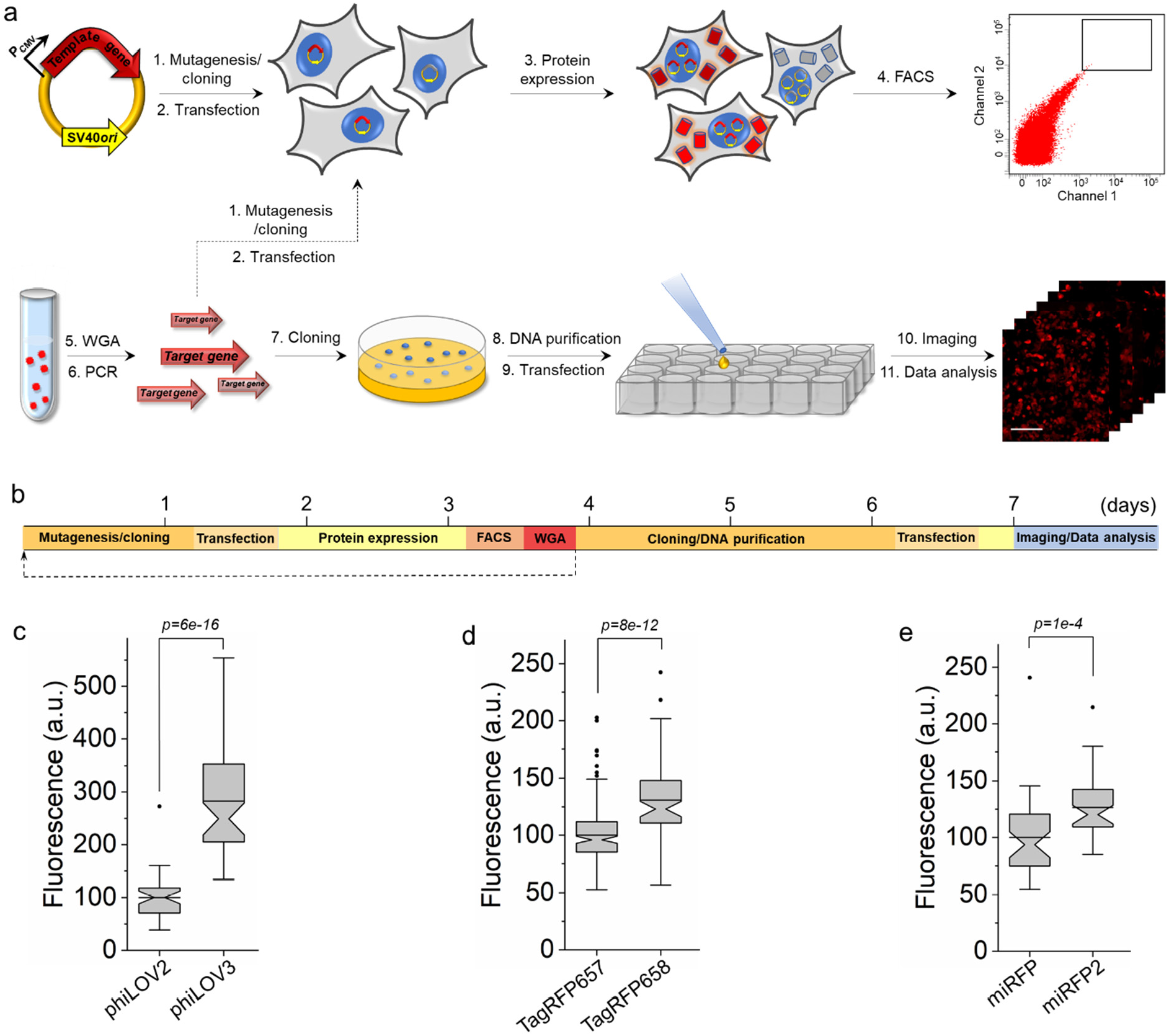
Rapid directed molecular evolution of fluorescent proteins in mammalian cells. (**a**) Workflow for directed evolution of proteins in HEK293FT cells using single-gene delivery via modified calcium phosphate transfection, fluorescence-activating cell sorting (FACS), and automated fluorescence imaging; WGA, whole-genome amplification. (**b**) Timeline of directed molecular evolution. (**c**) Green fluorescence of phiLOV2 and phiLOV3 expressed in HEK293FT cells (n= 40 and 59 cells, respectively, from two independent transfections for each protein; one-way ANOVA is used throughout this figure). Imaging conditions: excitation 475/34 nm from an LED, emission 527/50 nm. Box plots with notches are used throughout this paper, when n>9 (the narrow part of notch, median; top and bottom of the notch, 95% confidence interval for the median; top and bottom horizontal lines, 25% and 75% percentiles for the data; whiskers extend 1.5 × the interquartile range from the 25th and 75th percentiles; horizontal line, mean; dots represent outliers, data points which are less than the 25th percentile or greater than the 75th percentile by more than 1.5 times the interquartile range). (**d**) Near-infrared fluorescence of TagRFP657 and TagRFP658 expressed in HEK293FT cells (n= 181 and 76 cells, respectively, from two independent transfections for each protein). Imaging conditions: excitation 631/28 nm from an LED, emission 664LP. (**e**) Near-infrared fluorescence of miRFP and miRFP2 expressed in HEK293FT cells (n= 40 and 41 cells, respectively, from two independent transfections for each protein). Imaging conditions as in **d**.

### Spectral and biochemical characterization of the optimized FPs

To characterize spectral and biochemical properties of the selected proteins in comparison to their precursors, first, we expressed them in *E.coli* bacteria and purified using standard methods. The introduced mutations did not alter the spectral properties of the developed proteins when compared to the parental proteins. Absorption and fluorescence spectrum profiles of phiLOV3, TagRFP658, and miRFP2 appeared to be almost identical to that of their precursors with insignificant shifts in the maxima of the major bands (**Figure 2a-c**). Fluorescence excitation/emission peaks for phiLOV3 are at 452,477/502 nm, for TagRFP658 – at 611/658nm, and for miRFP2 – at 676/706 nm (**Table 1** and **Table 2**). The TagRFP658 and miRFP2 proteins exhibited similar intracellular photostability compared to the corresponding parental proteins, while phiLOV3 showed about 23% improvement in photostability halftime over its precursor (**Figure 2d-f**). The pH-stability of fluorescence for phiLOV3, TagRFP658, and miRFP2 was similar to that of the corresponding parental proteins and characterized by pK_a_ values of 3.3, 4.6, and 3.7, respectively (**Figure 2g-i, Table 1** and **Table 2**).

**Figure 2.**
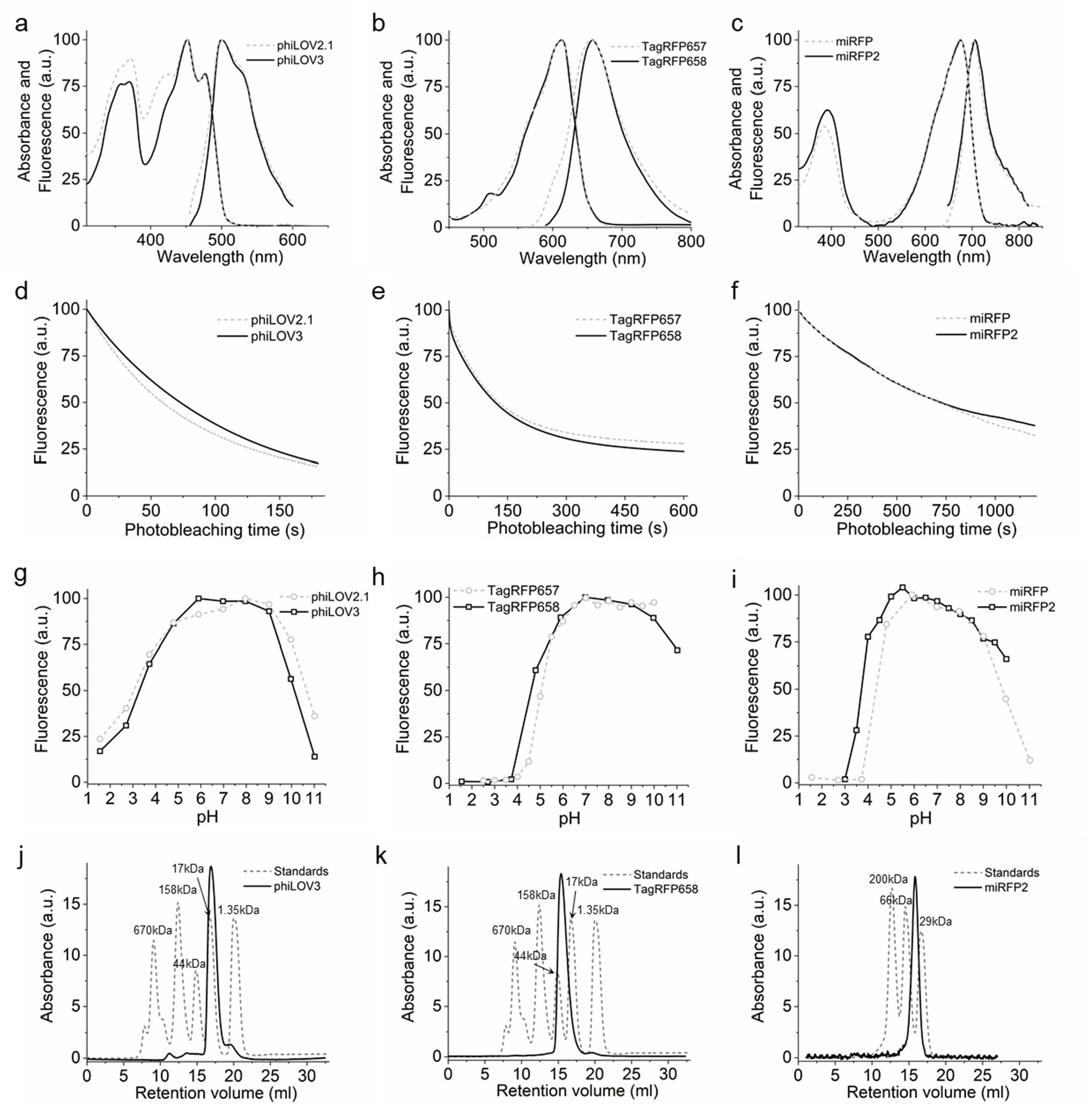
Spectroscopic and biochemical characterization of phiLOV3, TagRFP658, and miRFP2 in comparison to their precursors. (**a-c**) Absorbance and fluorescence emission spectra of (**a**) phiLOV2.1 (dashed line) and phiLOV3 (solid line), (**b**) TagRFP657 (dashed line) and TagRFP658 (solid line), (**c**) miRFP (dashed line) and miRFP2 (solid line). (**d**) Photobleaching curves of phiLOV2.1 (dashed line) and phiLOV3 (solid line) expressed in HEK293FT cells (n = 18 cells from 1 transfected sample, each). Photobleaching conditions: excitation 475/34 nm from an LED at ∼5 mW/mm^2^. (**e**) Photobleaching curves of TagRFP657 (dashed line) and TagRFP658 (solid line) expressed in HEK293FT cells (n = 16 cells from 1 transfected sample, each). Photobleaching conditions: excitation 628/31BP from a LED at 95 mW/mm^2^. (**c**) Photobleaching curves of miRFP (dashed line) and miRFP2 (solid line) expressed in primary cultured mouse neurons (n = 25 and 61 cells, respectively, from one culture, each). Photobleaching conditions: excitation 628/31BP from a LED at 88 mW/mm^2^. (**g-i**) pH dependence of fluorescence for (**g**) phiLOV2.1 (dashed line) and phiLOV3 (solid line), (**h**) TagRFP657 (dashed line; data from^32^) and TagRFP658 (solid line), and (**i**) miRFP (dashed line; data from^35^) and miRFP2 (solid line). (**j-l**) Size exclusion chromatography of (**j**) phiLOV3 at a concentration 3.8 mg/ml, (**k**) TagRFP658 at a concentration of 8 mg/ml, (**l**) miRFP at a concentration of 8.8 mg/ml (solid lines) and the indicated molecular weight standards (dashed lines).

**Table 1.** Properties of the FMN-binding phiLOV2.1 and phiLOV3 FPs.

| Protein | Abs. (nm) | Em. (nm) | EC (M<sup>-1</sup>cm<sup>-1</sup>) | QY (%) | Molecular brightness<sup>a</sup> | pK<sub>a</sub> | Brightness in HEK cells (%)<sup>b</sup> | Intracellular photo-stability (s)<sup>c</sup> |
| --- | --- | --- | --- | --- | --- | --- | --- | --- |
| phiLOV2.1 | 451, 476 | 501 | 13,500 | 20 | 2,700 | 3.0 | 100 | 59 |
| phiLOV3 | 452, 477 | 502 | 15,800 | 22 | 3,480 | 3.3 | 283 | 73 |

Abs – absorbance peak; Em – fluorescence emission peak; EC – extinction coefficient; QY – quantum yield; ND – not determined; <sup>a</sup>molecular brightness is a product of extinction coefficient and fluorescence emission quantum yield; <sup>b</sup>determined as mean green fluorescence relative to phiLOV2.1; <sup>c</sup>measured under continuous 470/25 nm wide-field illumination.

**Table 2.** Spectral and biochemical properties of the selected NIR FPs.

| Protein | Abs (nm) | Em (nm) | EC (M<sup>-1</sup>cm<sup>-1</sup>) | QY (%) | Molecular brightness<sup>a</sup> | pK<sub>a</sub> | Brightness in HEK cells (%)<sup>b</sup> | Photo-stability (s)<sup>c</sup> | Initial photo-bleaching rate (%/s)<sup>d</sup> | Ref |
| --- | --- | --- | --- | --- | --- | --- | --- | --- | --- | --- |
| mCardinal | 603 | 651 | 79,000 | 18 | 14,220 | 5.3 | 290 | 340 | 0.24 | 65 |
| mMaroon1 | 609 | 657 | 80,000 | 11 | 8,800 | 6.2 | 125<sup>e</sup> | ND | ND | 42 |
| iRFP670 | 643 | 670 | 114,000 | 11 | 15,540 | 4.0 | 154 | 245 | 0.56 | 38 |
| miRFP670nano | 645 | 670 | 95,000 | 10.8 | 10,260 | 3.7 | 8<sup>e</sup> | ND | ND | 19 |
| miRFP680 | 661 | 680 | 94,000 | 14.5 | 13,630 | 4.5 | 160 | ND | 0.06 | 20 |
| iRFP682 | 663 | 682 | 90,000 | 11 | 9,900 | 4.5 | 140 | 904 | 0.09 | 38 |
| miRFP703 | 674 | 703 | 90,900 | 8.6 | 7,820 | 4.5 | 48 | ND | ND | 66 |
| mRhubard713 | 690 | 713 | 113,500 | 7.6 | 8,630 | ND | 33 | ND | 0.02 | 39 |
| miRFP720 | 702 | 720 | 98,000 | 6.1 | 5,980 | 4.5 | 61 | ND | 0.03 | 40 |
| TagRFP657 | 611 | 657 | 34,000 | 10 | 3,400 | 5.0 | 61<sup>f</sup> | ND | ND | 32 |
| TagRFP658 | 611 | 658 | 41,200 | 10 | 4,120 | 4.6 | 79 | 640 | 0.12 | This study |
| miRFP | 674 | 703 | 92,400 | 9.7 | 8,960 | 4.3 | 79<sup>g</sup> | ND | ND | 67 |
| miRFP2 | 676 | 706 | 55,600 | 4.3 | 2,390 | 3.7 | 100 | 995 | 0.08 | This study |
| emiRFP2 | ND | ND | ND | ND | ND | ND | 205 | 1050 | 0.08 | This study |

Abs – absorbance peak; Em – fluorescence emission peak; EC – extinction coefficient; QY – quantum yield; ND – not determined; <sup>a</sup>molecular brightness is a product of extinction coefficient and fluorescence emission quantum yield; <sup>b</sup>determined as a Cy5-to-green fluorescence intensity ratio for live HEK cells relative to miRFP2 unless otherwise state; <sup>c</sup>measured in live HEK cells under continuous wide-field illumination with 631 nm laser at 66mW/mm<sup>2</sup>; <sup>d</sup>determined as a liner fit of the initial segment of photobleaching curve; <sup>e</sup>determined as mean Cy5 fluorescence intensity in live HEK cells relative to mCardinal; <sup>f</sup>determined as mean Cy5 fluorescence intensity in live HEK cells relative to TagRFP658; <sup>g</sup>determined as mean Cy5 fluorescence intensity in live HEK cells relative to miRFP2.

Previously we showed that EGFP and bacterial phytochrome photoreceptor (BphP)-derived FPs can be efficiently co-excited under two-photon excitation at 880 nm due to an overlap of their corresponding first S_0_→S_1_ and higher S_0_→S_n_^36^ electronic transitions. To explorer utility of TagRFP658 for two-photon microscopy, we measured its two-photon cross-sections in the range 790–1300 nm (**Supplementary Figure 6**). The two-photon spectrum had a major peak at 1230 nm coinciding with one-photon peak (at double wavelength) rather well and exhibited a strong feature in the region corresponding to the S_0_→S_n_ transitions with strong absorption only at wavelength below 830 nm. As a result, two-photon spectral overlap of TagRFP658 with EGFP is less substantial than that for iRFPs and might not be practical for single-wavelength co-excitation with EGFP. Oligomerization of FPs can result in increase in intracellular brightness by enabling a higher protein expression level. Therefore, we decided to determine the oligomeric state of the selected proteins. Size-exclusion chromatography demonstrated that the developed variants preserved monomeric state even at a high concentration (**Figure 2g-i**, see **Supplementary Figure 7** for calibration plots). The molecular brightness of phiLOV3 and TagRFP658 was about 29% and 21% higher than that of their precursors, respectively, while molecular brightness of miRFP2 was 3.7-fold lower than that of miRFP (**Table 1** and **Table 2**). Improvement of the TagRFP568 intracellular brightness corresponded to increase in molecular brightness over its precursor. However, relative intracellular brightness of phiLOV3 and miRFP2 was several folds higher than their molecular brightness when compared to phiLOV2.1 and miRFP, respectively (**Table 1** and **Table 2**). Perhaps this can be explained by the difference in binding affinities of the FMN (flavin mononucleotide) and BV co-factors between the selected proteins and their precursors.

Therefore, to investigate the significant difference in relative brightness of miRFP2 *in vitro* and in cell culture, we decided to estimate BV binding efficiency in HEK cells. We measured intracellular brightness of HEK cells expressing miRFP703, miRFP, or miRFP2 under different concentrations of exogenously administrated BV. The miRFP703 protein was used as an additional reference since it shares the highest amino acid identity (∼93%) with miRFP2 and was reported to have higher than miRFP2 molecular brightness^20,37^ (**Table 2**). Addition of BV at 62.5 µM resulted in 3.2- and 3.7-fold increase in intracellular brightness of miRFP and miRFP703 while miRFP2 showed only 1.8-fold increase (**Supplementary Figure 8**). The data suggested that miRFP2 has higher BV binding affinity and as a result larger fraction of miRFP2 exists in the BV-bound state in HEK cells. The higher BV affinity of miRFP2 can account for its higher intracellular brightness over miRFP and miRFP703, which however have superior molecular brightness when measured in solution using the proteins purified from *E.coli*. Overall, characterization *in vitro* and in cell culture showed that enhancement of intracellular brightness was not due to changes in spectral properties or oligomeric state of the proteins but rather due to improving either molecular brightness in case of TagRFP658 or cofactor binding affinity in case of miRFP2.

Next, we compared intracellular brightness of TagRFP658 and miRFP2 in NIH3T3 mouse embryonic fibroblasts and PAC2 zebrafish embryonic fibroblasts under identical imaging conditions. While finalizing this study, it was reported that swapping the N-terminus of the *Rp*BphP1-derived miRFPs with that of the *Rp*BphP2 protein can significantly improve intracellular brightness in mammalian cells without affecting molecular brightness^20^. Following the described strategy, we generated the enhanced version of miRFP2, or emiRFP2 for short, and used it for side-by-side comparison with TagRFP658 and miRFP2. To perform expression-level independent quantification of intracellular brightness, the NIR-FPs were co-expressed with mClover3 under the EF1α:2xCMV:EF1α bidirectional promoter (**Figure 3**). Under transient expression in NIH3T3 cells, emiRFP2 exhibited 1.4- and 4.8-fold higher fluorescence than TagRFP658 and miRFP2, respectively, when quantified using Cy5-to-green fluorescence ratio (**Figure 3b**; here and below we reported fluorescence intensities without normalization to excitation and emission efficiencies for spectrally distinct FPs, unless otherwise indicated, to provide direct comparison of intracellular brightness in real experimental settings rather than intrinsic properties of the FPs). Similar results were obtained in PAC2 fibroblasts with emiRFP2 being 1.5- and 4.1-fold brighter than TagRFP658 and miRFP2, respectively (**Figure 3d**). Administration of exogenous BV at 25 µM in PAC2 cells further increased fluorescence of miRFP2 and emiRFP2 by 2- and 2.3-fold, respectively, although the increase of miRFP2 brightness was not statistically significant (**Figure 3d**). Consistently with the previously reported results^20^, swapping the N-terminus of miRFP2 significantly improved its intracellular brightness. The emiRFP2 protein also outperformed TagRFP658 in terms of intracellular brightness in Cy5 channel.

**Figure 3.**
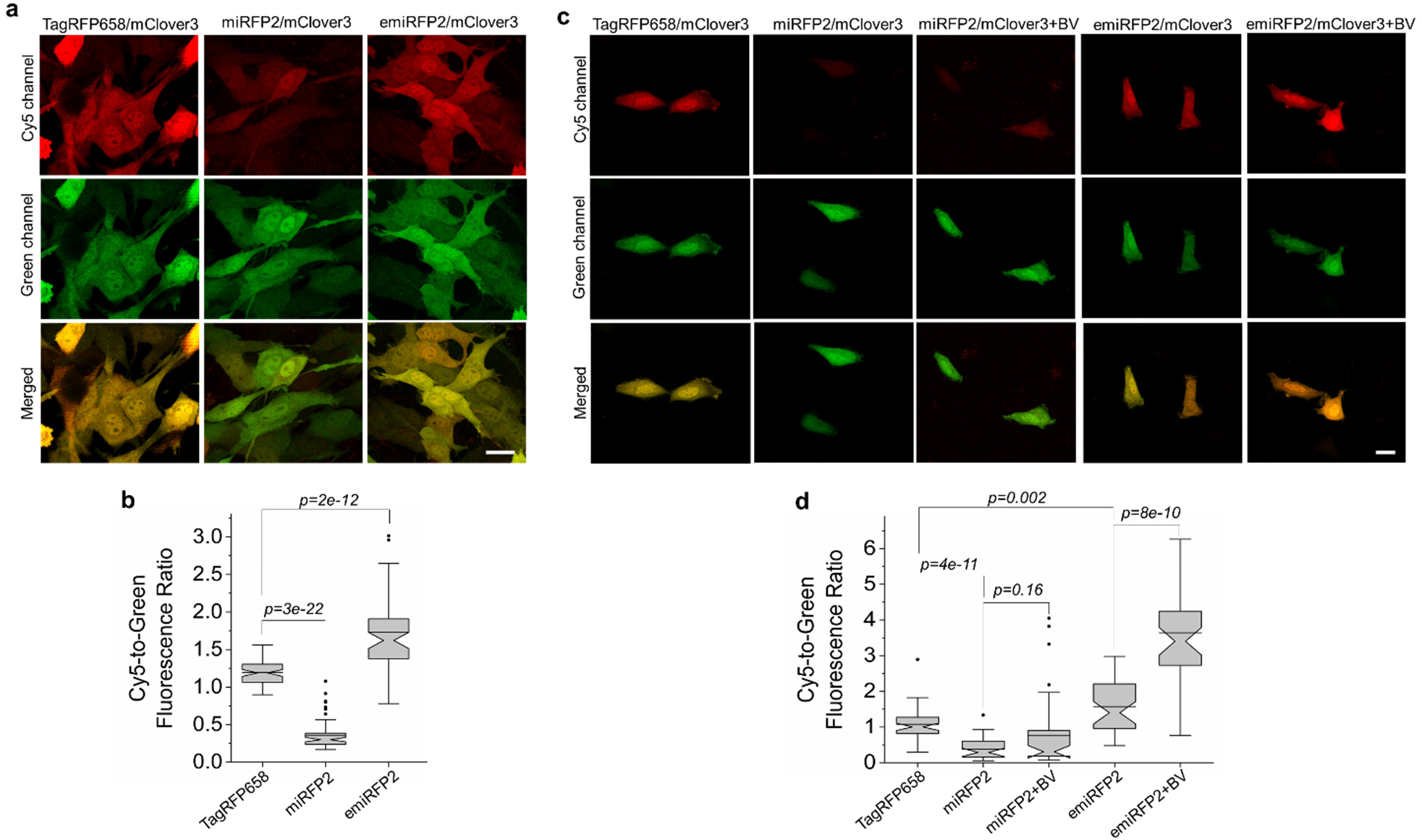
Fluorescence imaging of TagRFP658, miRFP2, and emiRFP2 expressed in mammalian and zebrafish cell cultures. (**a**) Representative fluorescence images of live NIH3T3 mouse embryonic fibroblasts co-expressing TagRFP658, miRFP2, and emiRFP2 with mClover3 under the EF1α:2xCMV:EF1α bidirectional promoter (n= 67, 62, and 59 cells, respectively, from one transfection each; the dynamic range of the images for the same channel was kept constant throughout the panels **b** and **d**). Imaging conditions: Cy5 channel, excitation 633 nm from a laser, emission 660-785 nm; green channel, excitation 488 nm from a laser, emission 495-530 nm). (**b**) Cy5-to-green fluorescence ratio of live NIH3T3 fibroblasts shown in **a** (n= 67, 62, and 59 cells for TagRFP658, miRFP2, and emiRFP2, respectively, from one transfection each; Kruskal–Wallis ANOVA). Box plots with notches are used in this figure (see Fig. 1c for the description). (**c**) Representative fluorescence images of live Pac2 zebrafish embryonic fibroblasts co-expressing TagRFP658, miRFP2, and emiRFP2 with mClover3 under the EF1α:2xCMV:EF1α bidirectional promoter with and without addition of 25 µM BV for 3 h before imaging (n= 49, 36, 41, 39 and 39 cells for TagRFP658, miRFP2, miRFP2+BV, emiRFP2, and emiRFP2+BV, respectively, from one transfection each). Imaging conditions as in a. (**d**) Cy5-to-green fluorescence ratio of live Pac2 fibroblasts shown in c with and without addition of 25 µM BV for 3 h before imaging (n= 49, 36, 41, 39 and 39 cells for TagRFP658, miRFP2, miRFP2+BV, emiRFP2, and emiRFP2+BV, respectively, from one transfection each; Kruskal–Wallis ANOVA). Imaging conditions as in **a**. Scale bars, 20 μm.

It is important to validate if the FPs optimized in mammalian cells outperform their counterparts evolved in a bacterial system. Therefore, we decided to compare intracellular brightness and photostability of TagRFP658, miRFP2, and emiRFP2 to spectrally similar GFP-like and BphP-based NIR-FPs in HEK cells using a standard Cy5 filter set with a wide bandpass emission filter (730/140 nm), which allows efficient collection of NIR fluorescence. Based on the literature search, we selected a set of monomeric and dimeric NIR FPs including mCardinal^11^, iRFP670 (*ref.*^38^), miRFP680 (*ref.*^20^), iRFP682 (*ref.*^38^), mRhubarb713 (*ref.*^39^), and miRFP720 (*ref.*^40^), which were reported to have high performance in cultured mammalian cells. To account for the expression level during transient transfection of HEK cells, the NIR FPs were co-expressed with EGFP using the self-cleavage P2A peptide^41^. First, we compared the intracellular brightness of the BphP-based NIR FPs independent of spectral properties. We left mCardinal and TagRFP658 out of this comparison because these two proteins belong to a different spectral class (∼30-40 nm spectral shift). To correct for the difference in the fluorescence spectra, raw fluorescence intensity values were normalized to absorbance of the proteins at the excitation wavelength and overlap of the fluorescence spectrum with the transmission of the emission filters and quantum efficiency of the sCMOS camera chip. Quantification of the normalized intracellular fluorescence revealed that miRFP2 was dimmer than iRFP670, miRFP680, and iRFP682 by 14%, 45%, and 27%, respectively, but brighter than mRhubarb713 and miRFP720 by 121% and 7%, respectively. In turn, emiRFP2 outperformed all other BphP-based NIR FPs exhibiting 80%, 40%, 61%, 353%, and 120% higher normalized fluorescence than iRFP670, miRFP680, iRFP682, mRhubard713, and miRFP720, respectively. To evaluate the applicability of the NIR FPs for imaging in the Cy5 channel, we compared expression level-normalized fluorescence and intracellular photostability. mCardinal exhibited the highest intracellular fluorescence followed by emiRFP2, and TagRFP658 was 3.6- and 2.6-fold dimmer than mCardinal and emiRFP2, respectively (**Figure 4a**, see **Table 2** for the quantification of all proteins). It should be noted that the expression level of the FPs as assessed by fluorescence intensity of GFP was comparable for all expressed constructs. Next, we assessed intracellular photostability under continuous wide-field illumination at 66 mW/mm^2^ light density, which was about 2-5-fold higher than we routinely used for NIR-FPs imaging in mammalian cell culture. Higher light power was used in order to achieve faster photobleaching and speed up imaging process. Photostability of TagRFP658, miRFP2, and emiRFP2 was ∼2, 3, and 3-fold higher than that of mCardinal, respectively, however, lower than that for miRFP680, mRhubard, and miRFP720 (**Figure 4b** and **Table 2**). Based on the assessed characteristics, emiRFP2 and miRFP680 exhibited the best combination of brightness and photostability among the tested NIR FPs. However, as for the majority of in vivo applications, intracellular brightness is usually the most crucial property we decided to use mCardinal as a major reference for further characterization of the developed NIR FPs in neurons. Moreover, mCardinal outperformed other recently published NIR FPs, such as mMaroon^42^, and miRFP670nano^19^, when compared side-by-side in HEK cells (**Supplementary Figure 9**).

**Figure 4.**
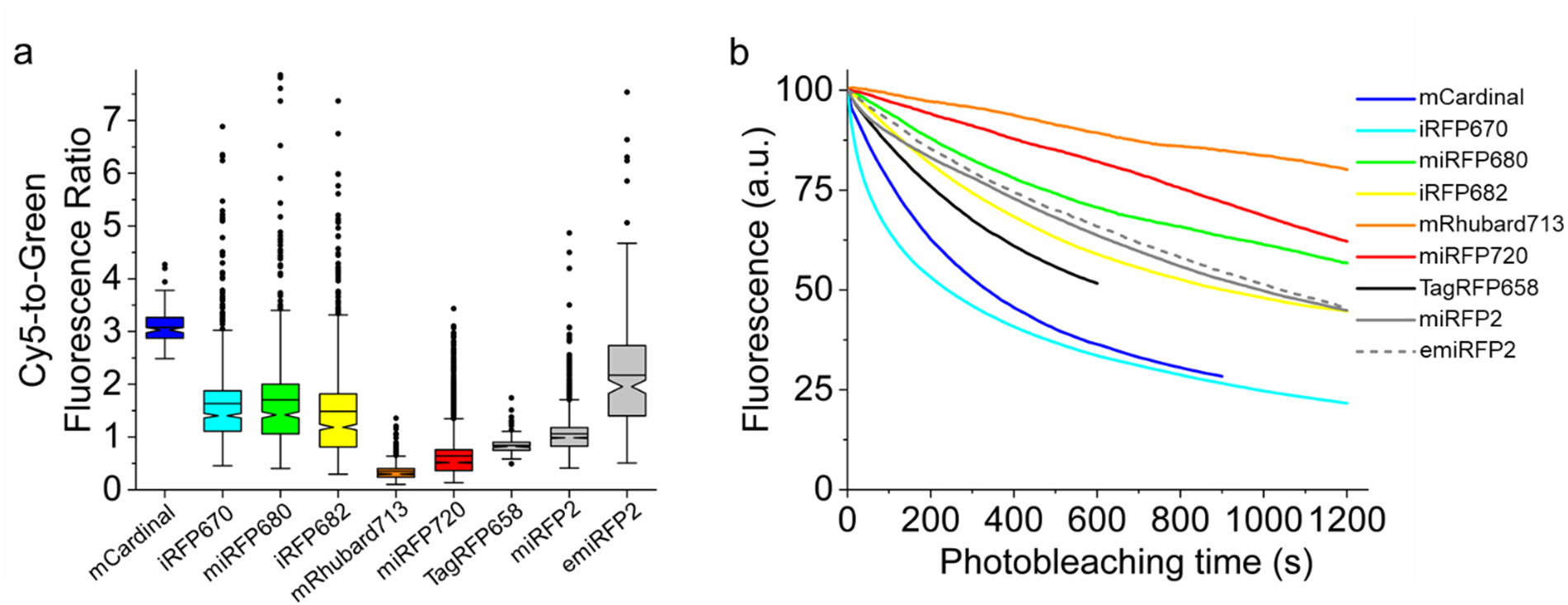
Intracellular brightness and photostability of the selected NIR FPs in live HEK cells. (**a**) Cy5-to-green fluorescence ratio for NIR-FPs co-expressed with GFP via the P2A peptide (n = 144, 775, 612, 837, 446, 2937, 226, 1981, and 271 cells for mCardinal, iRFP670, miRFP680, iRFP682, mRhubarb713, miRFP720, TagRFP658, miRFP2, and emiRFP2, respectively, from 2 independent transfections each). Box plots with notches are used in this figure (see Figure 1c for the full description). Imaging conditions: Cy5 fluorescence, excitation 635/22 nm from 637 nm laser, emission 730/140 nm; green fluorescence, excitation 478/24 nm for an LED; emission 535/46 nm. (**b**) Photobleaching curves for NIR-FPs under continuous wide-field illumination from 637 nm laser at 66 mW/mm^2^ (n = 55, 32, 10, 43, 32, 48, 56, 38, and 37 cells for mCardinal, iRFP670, miRFP680, iRFP682, mRhubarb713, miRFP720, TagRFP658, miRFP2, and emiRFP2, respectively, from 2 independent transfections each).

### TagRFP658 as a bright and photostable near-infrared fluorescence tag for neuronal labeling

TagRFP658, along with its precursor TagRFP657, possesses the most red-shifted fluorescence excitation maximum (611 nm) among all monomeric GFP-like FPs that enables efficient excitation with red light excitation sources available on the common imaging systems. Therefore, we decided to explore the utilization of TagRFP658 as a NIR fluorescent probe for neuronal labeling under wide-field and light-sheet microscopy using standard Cy5 filter set. As a reference, we used spectrally similar GFP-like NIR FP, mCardinal, which exhibited the highest intracellular brightness among tested NIR-FPs (**Table 2**). First, we cloned the mCardinal and TagRFP658 genes under CaMKII promoter and expressed them in primary hippocampal mouse neurons using calcium phosphate transfection. The fluorescence of TagRFP658 was evenly distributed within the cytosol, individual dendrites, and nucleus of live cultured neurons without any aggregation or nonspecific localization even during long-term expression for up to 23 days in vitro (DIV; **Figure 5a,b**). Quantification of the intracellular brightness under red light excitation at 628/14 nm showed that TagRFP658 is about 28% brighter than mCardinal although not statistically significant (**Figure 5c**). In addition, intracellular photostability of TagRFP658 under continuous wide-field excitation was almost twice higher than that of mCardinal (photobleaching half-time for TagRFP658 and mCardinal was 164 s and 80 s, respectively; **Figure 5d**). Using whole-cell patch clamp recordings, we showed that TagRFP658 expression did not alter membrane resistance, membrane capacitance, or the resting potential of cultured mouse neurons (**Supplementary Figure 10**).

**Figure 5.**
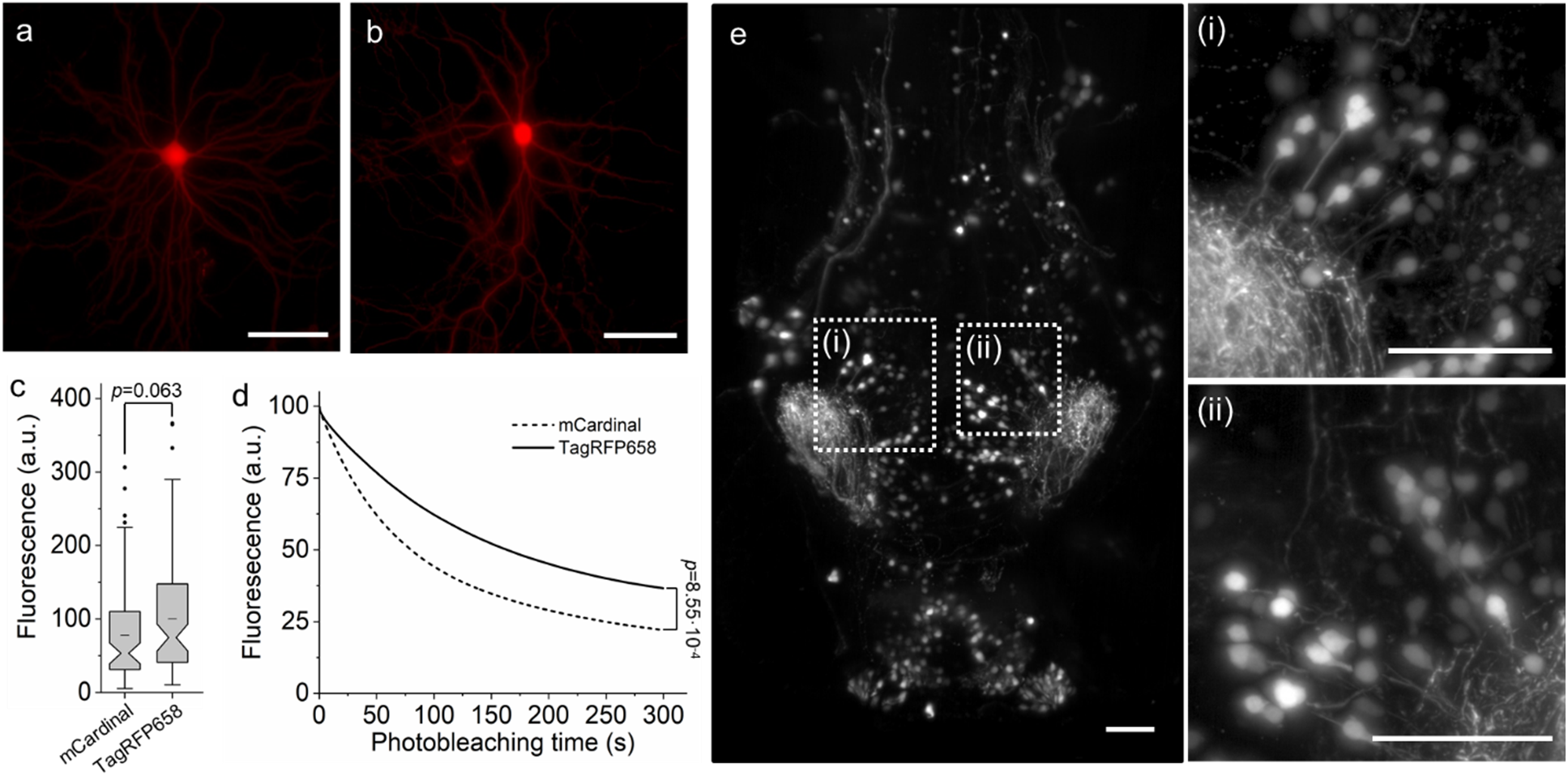
Characterization of TagRFP658 in neurons in primary culture and *in vivo*. (**a,b**) Representative fluorescence images of primary cultured mouse hippocampal neurons expressing TagRFP658 at (**a**) 14 and (**b**) 23 days in vitro (DIV; n = 53 and 33 neurons, respectively, from two independent cultures). Imaging condition: excitation 631/28 nm from an LED, emission 664LP. (**c**) Relative fluorescence of cultured mouse hippocampal neurons expressing mCardinal and TagRFP658 (n = 78 and 85 neurons, respectively, from two independent cultures for each protein; one-way ANOVA). Imaging conditions as in **a**. (**d**) Raw photobleaching curves for mCardinal (dashed line) and TagRFP658 (solid line) in primary cultured mouse hippocampal neurons (n = 9 and 7 neurons, respectively, from one culture each; one-way ANOVA). Imaging condition: excitation 631/28 nm from an LED at 70 mW/mm^2^, emission 664LP. (**e**) Representative light sheet image of head of zebrafish larvae at 4 day post fertilization expressing TagRFP658 in neurons (n=10 fish from two independent injections). Imaging conditions: excitation 638 nm from a laser, emission 665LP. (**i, ii**) High-magnification images of the respective regions shown in white boxes in **e**. Scale bars, 50 μm.

We next explored the use of TagRFP658 as a neuronal fluorescent label in zebrafish larvae (*Danio rerio*) under light sheet microscopy. We transiently expressed a zebrafish codon-optimized version of TagRFP658 in a subset of neurons and visualized fluorescence in live zebrafish larva at 5 days post fertilization. The TagRFP658 protein exhibited well-detectable near-infrared fluorescence, which was evenly distributed in cell bodies and processes (**Figure 5e**). High photostability and intracellular brightness make TagRFP658 as a promising NIR tag for imaging neurons in culture and *in vivo*.

### miRFP2 as a near-infrared fluorescence tag for neuronal labeling

Despite of more than 50 nm shift in fluorescence spectra between TagRFP658 and miRFP2, both proteins can be efficiently visualized using a Cy5 filter set with broad band pass or long pass emission filter (e.g., see **Figure 3**), which is commonly available on the standard imaging systems. However, a Cy5.5 filter set may fit fluorescence spectrum of miRFP2 better than Cy5 filter due to slightly red shifted wavelength. To compare efficiency of fluorescence imaging in Cy5 and Cy5.5 channels, we equipped a wide-field microscope with commercially available 680-nm laser and used 710 nm long pass filter to collect fluorescence emission. For assessing intracellular characteristics in neurons, we selected mCardinal, TagRFP658, and emiRFP2. Since emiRFP2 outperformed miRFP2 in earlier experiments it was not used for quantitative imaging in neurons, although miRFP2 can be efficiently expressed and imaged in cultured neurons both under transient transfection and rAAV transduction (**Supplementary Figure 11**). The NIR FPs were co-expressed with GFP via P2A peptide under the CAG promoter in primary cultured neurons and imaged in Cy5, Cy5.5, and GFP channels. To perform fair comparison of fluorescence intensity in Cy5 and Cy5.5 channels, we acquired images under matching excitation power (66 mW/mm^2^) and the same exposure time (100 ms). The fluorescence of the NIR FPs was evenly distributed within the cytosol, individual dendrites, and nucleus of live cultured neurons without any aggregation or nonspecific localization (**Figure 6a**). Quantification of fluorescence intensity revealed that emiRFP2 were about 3.5-times brighter in Cy5.5 channel than in Cy5 channel, while the mCardinal fluorescence in Cy5.5 channel was almost undetectable (**Figure 6b**). When quantified by Cy5-to-green fluorescence ratio, mCardinal was 2.2- and 1.5-fold brighter than TagRFP658 and emiRFP2, respectively. However, when raw mean fluorescence values in Cy5 channel were compared, TagRFP658 was about 20% brighter than mCardinal, similarly to the results shown in **Figure 5c** (note comparable fluorescence intensity for the representative images of mCardinal and TagRFP658 in Cy5 channel, but significantly dimmer GFP fluorescence in case of mCardinal-P2A-GFP construct). Intracellular photostability of emiRFP2 was 4-fold lower under Cy5.5 illumination compared to Cy5 excitation (photobleaching half-times were 230 and 990 seconds under Cy5.5 and Cy5 illumination, respectively; **Figure 6c**). At the same time, emiRFP2 exhibited superior intracellular photostability compared to mCardinal and TagRFP658 in the Cy5 channel, which were characterized by photobleaching halftime of 340 and 700 seconds, respectively, closely matching corresponding values obtained in HEK cells (**Figure 6c** and **Table 2**). Similar correlations of intracellular brightness and photostability in Cy5 and Cy5.5 channels were also observed in live HEK cells (**Supplementary Figure 12**). These results demonstrated that Cy5.5 channel provided high efficiency of the emiRFP2 fluorescence imaging however the gain in brightness came at the cost of reduced photostability.

**Figure 6.**
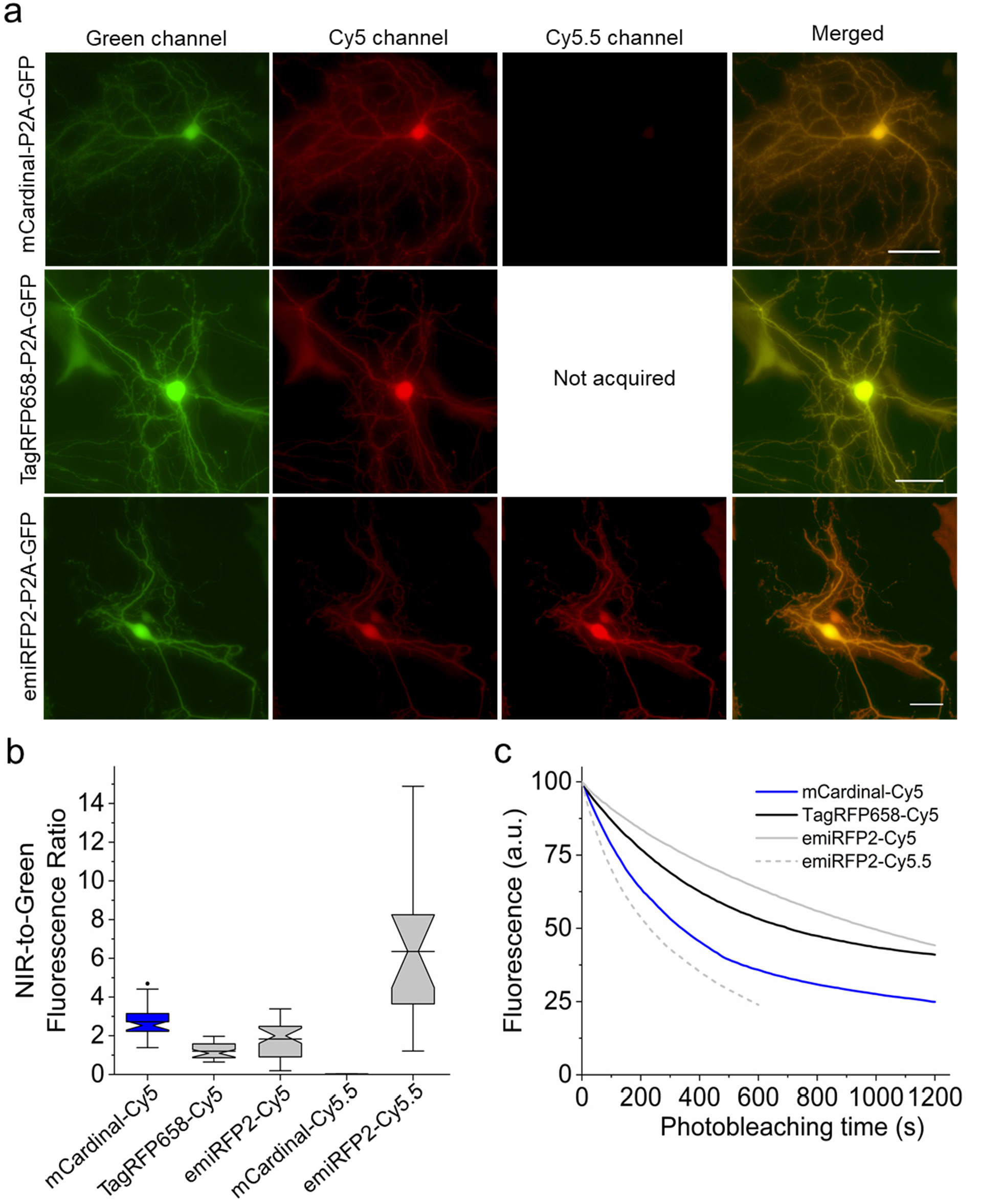
Intracellular brightness and photostability of mCardinal, TagRFP658, and emiRFP2 in live cultured hippocampal mouse neurons. (**a**) Representative fluorescence images of cells transfected with pAAV-CAG-mCardinal-P2A-GFP (top), pAAV-TagRFP658-P2A-GFP (middle), and pAAV-emiRFP2-P2A-GFP (n = 39, 33, and 41 neurons from 3, 2, and 3 independent transfections from one culture each for mCardinal, TagRFP658, and emiRFP2, respectively, for Cy5 channel and n = 15 and neurons from one independent transfection from one culture each for mCardinal and emiRFP2, respectively, for Cy5.5 channel). Imaging conditions: Cy5 channel: excitation 635/22 nm from 637 nm laser, emission 730/140 nm; Cy5.5 channel: excitation 680/13 nm from 680 nm laser, emission 710 LP; GFP channel: excitation 478/24 nm for an LED; emission 535/46 nm. Images in Cy5 and Cy5.5 were taken under matching excitation intensity (66 mW/mm^2^) and the same exposure time (100 ms). The dynamic range of fluorescence intensity in Cy5 and Cy5.5 channels are identical across all images. Scale bar, 20 µm. (**b**) NIR-to-green fluorescence ratio for mCardinal, TagRFP658, and emiRFP2 for the experiment shown in a. Box plots with notches are used in this figure (see Figure 1c for the full description). (**c**) Intracellular photostability of mCardinal, TagRFP658, and emiRFP2 in Cy5 and Cy5.5 channels (n = 8, 7, and 9 neurons from 3, 2, and 3 independent transfections from one culture each for mCardinal, TagRFP658, and emiRFP2, respectively, under Cy5 excitation and n = 5 neurons from one transfection from one culture for emiRFP2 under Cy5.5 excitation). Imaging conditions the same as in a.

High performance of miRFP2 and emiRFP2 in cultured neurons encouraged us to express them *in vivo* in model organisms such as mice, *C.elegans*, and *Drosophila melanogaster*. Since Cy5.5 filter set is not available on the majority of common imaging setups, all further characterization and validation of miRFP2 and emiRFP2 *in vivo* were performed using Cy5 filter set. First, we co-expressed mCardinal and emiRFP2 with GFP in cortical neurons in mice via *in utero electroporation* and performed imaging of the expressed proteins in acute brain slices at around P28. The emiRFP2 protein expressed in vivo showed even distribution in cell bodies and processes without aggregation (**Figure 7a**). Quantitative imaging showed that mCardinal had 4.8-fold higher Cy5-to-green fluorescence ratio than emiRFP2, however mean fluorescence intensity of mCardinal was only 1.7-fold higher than that of emiRFP2 (**Figure 7b,c**; note significantly lower green fluorescence intensity in the representative image for CAG-mCardinal-P2A-GFP construct compared to CAG-emiRFP2-P2A-GFP in **Figure 7a**). In addition, values for Cy5-to-green fluorescence ratio in the case of emiRFP2 exhibited significant variability ranging from 0.07 to 4.2 vs only 1.1 to 2.5 for mCardinal. The ratio variability can be visualized in the merged image of emiRFP2-P2A-GFP expressing neurons representing reddish, yellowish, and greenish neurons as a result of different levels of the emiRFP2 and GFP expression (compare to more even color distribution for mCardinal-P2A-GFP expressing neurons; **Figure 7a**). Next, we expressed the codon-optimized genes of miRFP2 using pan-neuronal expression systems in transgenic *C.elegans* and in *Drosophila* fruit flies. In case of *C.elegans* we did not observe any specific NIR signal, while NIR fluorescence in larvae and adult fruit flies was clearly detectable although its intensity was several times lower than in cultured neurons under the same imaging conditions (**Supplementary Figure 13**). Reduced fluorescence of miRFP2 can be due to the insufficient concentration of the BV cofactor in worms and flies. Previously it was shown that co-expression of heme oxygenase-1 (HO1) in worms and flies can enable fluorescence of the BphP-based FPs^43^ and biosensors^44^. To optimize conditions for miRFP2 expression we constructed two bicistronic vectors using IRES2 and a viral 2A cleavage sequence to co-express miRFP2 and HO1 and transfected them into HEK cells. Quantification of NIR fluorescence in HEK cells revealed that HO1 co-expression via IRES2 and P2A improves miRFP2 brightness by 40% and 83%, respectively (**Supplementary Figure 14**). Therefore, we decided to use the viral 2A cleavage sequence to co-express codon-optimized genes of miRFP2 and HO1 in neurons in worms and flies. Indeed, co-expression of HO1 enabled bright NIR fluorescence of miRFP2 in *C.elegans* and in larvae and adult fruit flies allowing visualization of individual neurons (**Figure 7d-g**). To validate miRFP2 performance in fruit flies we decided to compare its fluorescence to that of other GFP-like and BphP-derived NIR FPs under identical expression conditions. We establish transgenic lines with pan-neuronal expression of *Drosophila* codon optimized genes of mCardinal, mMaroon, iRFP-VC (aka iRFP713/V256C), and iRFP-VC-P2A-HO1 and quantified their Cy5 fluorescence in live intact 3^rd^ instar larva and adult fruit flies (**Figure 7h,i**). Co-expression of HO1 significantly enhanced brightness of miRFP2 and iRFP-VC both in adult flies and in larvae, although increase of miRFP2 brightness in adult flies was less pronounced than that in larva, 2.6-fold vs 9-fold. In the case of iRFP-VC, fluorescence enhancement in larva and adult flies was comparable, about 12- and 8-fold respectively. While miRFP2-T2A-HO1 outperformed mCardinal and mMaroon in terms of brightness in 3^rd^ instar larva, it was noticeably dimmer in adult fruit flies (**Figure 7h,i**). At the same time co-expression of iRFP-VC with HO1 resulted in the brightest fluorescence among the tested proteins both in larva and adult flies. Thus, we demonstrated that miRFP2 is a suitable NIR FP for imaging neurons in culture and *in vivo* in small model organisms, like worms and flies, and outperformed other high performing NIR FPs under certain conditions, e.g., in fruit fly larva. Our data also showed that co-expression of HO1 in worms and flies are essential for achieving sufficient brightness of BphP-based NIR FPs in *C.elegans* and *Drosophila* flies.

**Figure 7.**
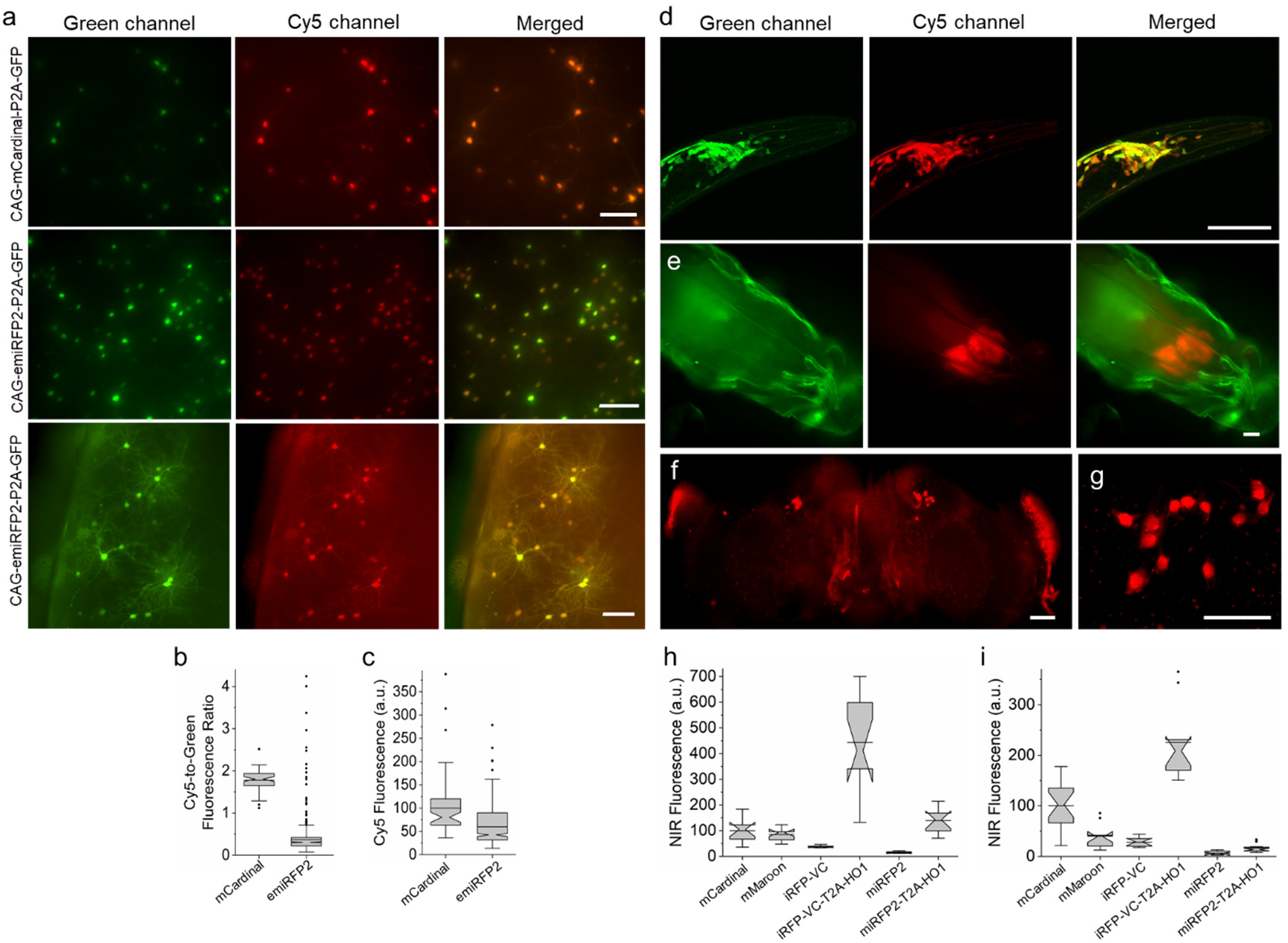
*In vivo* imaging and characterization of miRFP2 and emiRFP2. (**a**) Representative fluorescence images of live brain slices mCardinal-P2A-GFP (top) and emiRFP2-P2A-GFP (middle and bottom; n = 6 slices from 3 mice each). Imaging conditions: green channel: excitation 478/24 nm for an LED, emission 535/46 nm; Cy5 channel: excitation 635/22 nm from 637 nm laser, emission 730/140 nm. Top and middle rows of images are presented using the same dynamic range to facilitate visual comparison of mCardinal and emiRFP2 expressing neurons, bottom row images have adjusted LUTs to visualize processes of the emiRFP2 positive neurons. Scale bars, 100 µm. (**b**) Cy5-to-green fluorescence ratio of live neurons shown in **a** (n = 67 and 803 neurons from 3 mice each for mCardinal and emiRFP2, respectively). Imaging conditions same as in **a**. (**c**) Cy5 mean fluorescence of live neurons shown in **a** (n = 67 and 803 neurons from 3 mice each for mCardinal and emiRFP2, respectively). Imaging conditions same as in **a**. (**d**) Representative fluorescence images of the *C.elegans* head co-expressing codon-optimized genes of miRFP2-T2A-HO1 and jGCaMP7b in neurons (n=15 worms from two independent microinjections). Imaging conditions: green channel, excitation 488 nm from a laser, emission 500-650 nm; Cy5 channel, excitation 631 nm from a laser, emission 645-700 nm. Scale bar, 50 µm. (**e**) Representative fluorescence images of live intact 3^rd^ instar *Drosophila* larva expressing miRFP2-T2A-HO1 (n= 10 larvae from two transgenic lines). Imaging conditions: green channel, excitation 475/34 nm from LED, emission 527/50 nm (green fluorescence corresponds to autofluorescence); Cy5 channel, excitation 631/28 nm from LED, emission 665LP. Scale bar, 100 µm. (**f**) Representative low-magnification fluorescence image of brain explant from adult *Drosophila* fly expressing codon-optimized gene of miRFP2-T2A-HO1 in R84C10-GAL4 line (n= 5 brains from one transgenic lines). Imaging conditions: excitation 635/22 nm from 637 nm laser, emission 665LP. Scale bar, 50 µm. (**g**) Representative high-magnification fluorescence image of brain explant from adult *Drosophila* fly expressing codon-optimized miRFP2-T2A-HO1 in R84C10-GAL4 line (n= 5 brains from one transgenic lines). Imaging conditions: excitation 631 nm from a laser, emission 645-700 nm. Scale bar, 50 µm. (**h**) Relative NIR fluorescence of mCardinal, mMaroon, iRFP-VC, iRFP-VC-T2A-HO1, miRFP2, and miRFP2-T2A-HO1 expressed pan-neuronally in *Drosophila* 3^rd^ instar larvae (n=10, 12, 7, 11, 9, and 10 larvae from one transgenic line each, respectively). Imaging conditions same as in **g**. (**i**) Relative Cy5 fluorescence of mCardinal, mMaroon, iRFP-VC, iRFP-VC-T2A-HO1, miRFP2, and miRFP2-T2A-HO1 expressed pan-neuronally in adult *Drosophila* fly (n=19, 10, 11, 11, 11, and 21 flies from one transgenic line each, respectively). Imaging conditions same as in **g**.

### Multicolor imaging in cell culture and *in vivo*

Fluorescence spectra of TagRFP658 and miRFP2 are well separated from that of other commonly used GFPs and RFPs and therefore can enable multicolor crosstalk-free imaging using standard imaging filters. We decided to test the feasibility of TagRFP658 and miRFP2 in multicolor neuronal imaging in combination with green and red FPs expressed in larval zebrafish cerebellar Purkinje cells (PCs). First, we generated a set of the bidirectional expression constructs (**Figure. 3)** using zebrafish codon-optimized genes of NIR FPs allowing to co-express a nuclear-localized H2B histone fused to either mCardinal, TagRFP658, miRFP2 or emiRFP2 with cytoplasmic green FP mClover3 and transfected them into NIH3T3 cells stably expressing trans-Golgi network protein 46 (TGN46) fused to red FP mScarlet. Imaging using standard filter configuration under confocal microscope allowed for crosstalk-free triple color imaging for each combination of the selected FPs (**Supplementary Figure 15**). Next, we expressed the selected FPs in specific neuronal subpopulations of zebrafish larvae, especially targeting to cerebellar Purkinje cells (PCs) using the same constructs but carrying a PC specific bidirectional promoter (*ref.*^45^) instead of the ubiquitous CMV-EF1 promoter. As the enhanced variant of miRFP2, emiRFP2, exhibited higher intracellular brightness both in NIH3T3 cells and zebrafish PAC2 fibroblast than that of the original miRFP2 (**Figure. 3e-h**), it was selected for expression in zebrafish larva. Each construct was injected into the stable transgenic zebrafish embryos inducing PC specific expression of membrane tagged mScarlet at 4 dpf-larval zebrafish. Using a standard confocal microscope equipped with 488, 561, and 633 nm lasers we were able to easily visualize NIR fluorescence in PC nucleus using Cy5 channel together with mClover3 distributed throughout entire PC’s cytoplasm using green channel, while imaging in red channel provided clear visualization of the PCs axonal and/or dendritic structures with membrane tagged mScarlet fluorescence whose expressions are predominantly detected in PCs together with slight expression in tectal neurons (**Figure 8b,c**). Thus, NIR fluorescence of the tested NIR FPs were easily distinguishable from mScarlet fluorescence and thus can be used to label multiple neuronal compartments in conjunction with additional blue, or green fluorescent proteins. We further quantified the brightness and photostability of mCardinal, TagRFP658, and emiRFP2 fluorescence by expressing corresponding constructs in PCs of less pigmented brass embryos (**Supplementary Figure 16**). To account for expression level of the FPs, we calculated NIR-to-green fluorescence ratio for single PCs in zebrafish embryos. Fluorescence quantification revealed that mCardinal fluorescence was about twice higher than that of TagRFP658, whereas emiRFP2 exhibited 3.6-fold lower brightness than mCardinal (**Figure 8d**). Photobleaching experiments performed under identical excitation power for all selected FPs demonstrated 10-times higher photostability of TagRFP658 over mCardinal. However, emiRFP2 fluorescence exhibited rapid decay with half-time about 3 s making emiRFP2 about 10-fold less photostable than mCardinal at least in this zebrafish model (**Figure 8e**). Despite lower intracellular brightness of TagRFP658 than mCardinal, it can be more practical fluorescent tag for live imaging due to significantly higher photostability.

**Figure 8.**
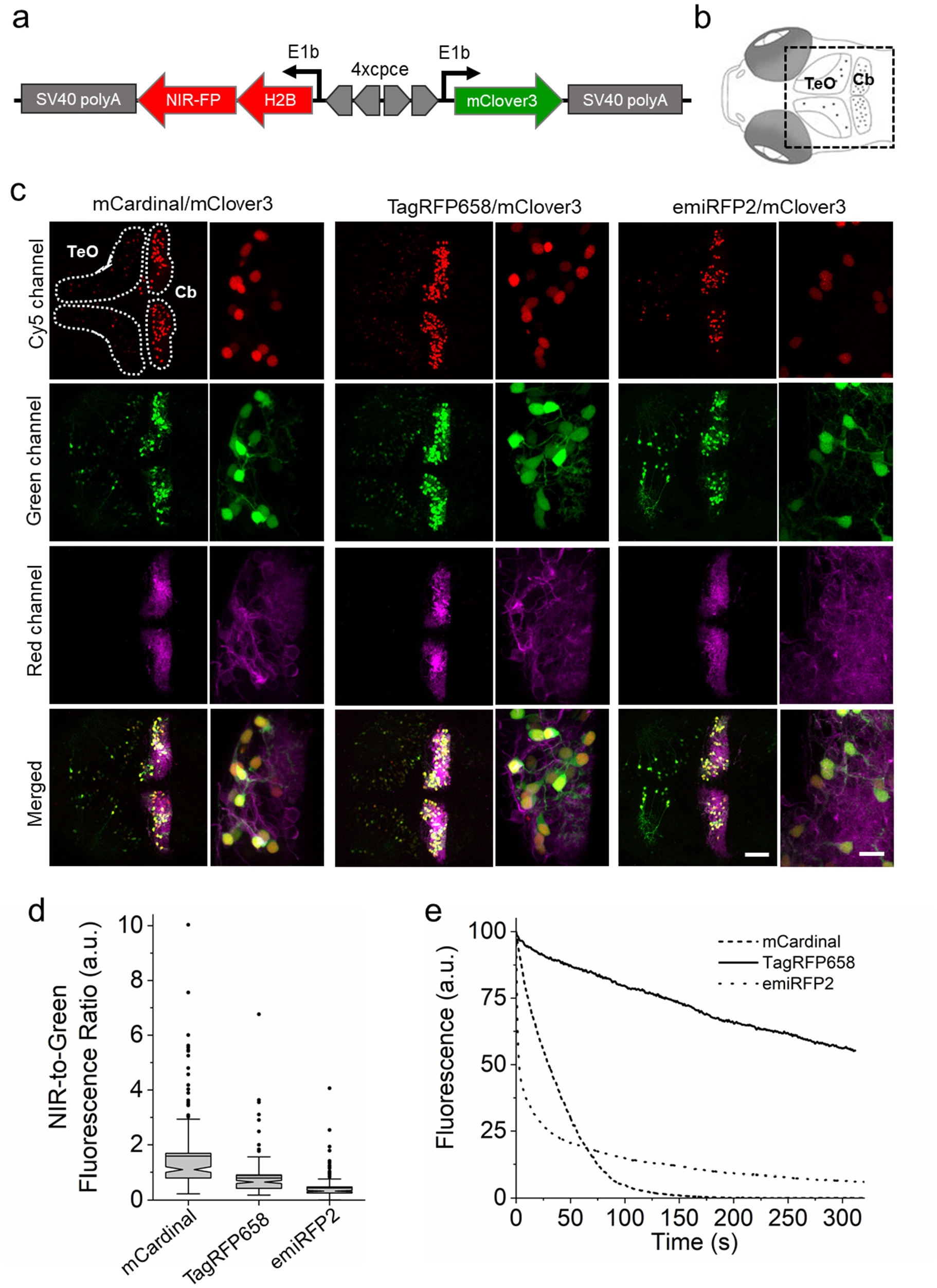
Three-color *in vivo* imaging of zebrafish larvae. (**a**) Schematic drawing of a bidirectional PC-specific expression system using 4x PC specific regulatory element (cpce) to co-express NIR-FP-H2B fusion protein with mClover3. (**b**) Schematic drawings of dorsal view of 4 dpf larval zebrafish brain region delineating the optic tectum (TeO) and the cerebellum (Cb). The region of interest enclosed by a square was recorded in images shown below. (**c**) Representative confocal images of 4 dpf zebrafish larvae transiently co-expressing H2B-NIR-FP with mClover3 (note: expression of this construct occurs in a mosaic manner) in stable transgenic background with PC specific membrane targeted mScarlet. The optic tectum (TeO) and the cerebellar region (Cb) are enclosed by the white dashed lines. Each subset of images (left, whole overview of tecum and cerebellar region; right, higher magnified images depicting PCs) shows the expression of each H2B-NIR-FP fusion (left, mCardinal; middle, TagRFP658; right, emiRFP2), cytoplasmic mClover3, membrane targeted mScarlet, and the overall merged image (upper to lower; n=4 fish for each subset). Imaging conditions, NIR channel: excitation 633 nm laser, emission 722/63 nm; green channel: excitation 488 nm laser, emission 513/17 nm; red channel: excitation 561 nm laser, emission 585/15 nm. Scale bars: 50 µm (overviewed images), 10 µm (higher magnified images) (**d**) NIR-to-green fluorescence ratio for NIR-FP expressing PCs (n=175, 161, 192 cells for mCardinal, TagRFP658, and emiRFP2, respectively, from 4 fish each; imaging conditions as in **c**). Box plots with notches are used (see caption for Figure 1c for the full description). (**e**) Photostability curves for mCardinal (dashed line), TagRFP658 (solid line), and emiRFP (dotted line) expressed in PCs (n=4 cells for each from one fish each).

Earlier we demonstrated that (e)miRFP2 imaging in Cy5.5 channel is several times more efficient than in Cy5 channel, while mCardinal fluorescence is not detectable in Cy5.5 channel (**Figure 6a,b** and **Supplementary Figure 12**). Therefore, we decided to explore a possibility for dual-color near-infrared imaging of subcellular structures using combination blue-shifted and red-shift NIR FPs, *e.g.*, mCardinal and emiRFP2. First, we verified that the developed NIR FPs can be properly localized in fusions with structured proteins in mammalian cells. Indeed, fusions of TagRFP658 and (e)miRFP2 with α-tubulin, β-actin, and keratin demonstrated proper localization in cultured mammalian cells (**Figure 9a-f**). Next, we co-expressed Mito-mCardinal and H2B-emiRFP2 fusions in HeLa cells and acquired images in Cy5 and Cy5.5 channels. To further improve spectral separation, we swapped wide bandpass emission filter in our standard Cy5 channel with narrower emission filter 679/41 nm. This optical setup enabled crosstalk free imaging of mCradinal and emiRFP2 (**Figure 9g**). The blue-shifted and red-shifted NIR FPs can further increase spectral multiplexing of fluorescence imaging in combination with other common FPs.

**Figure 9.**
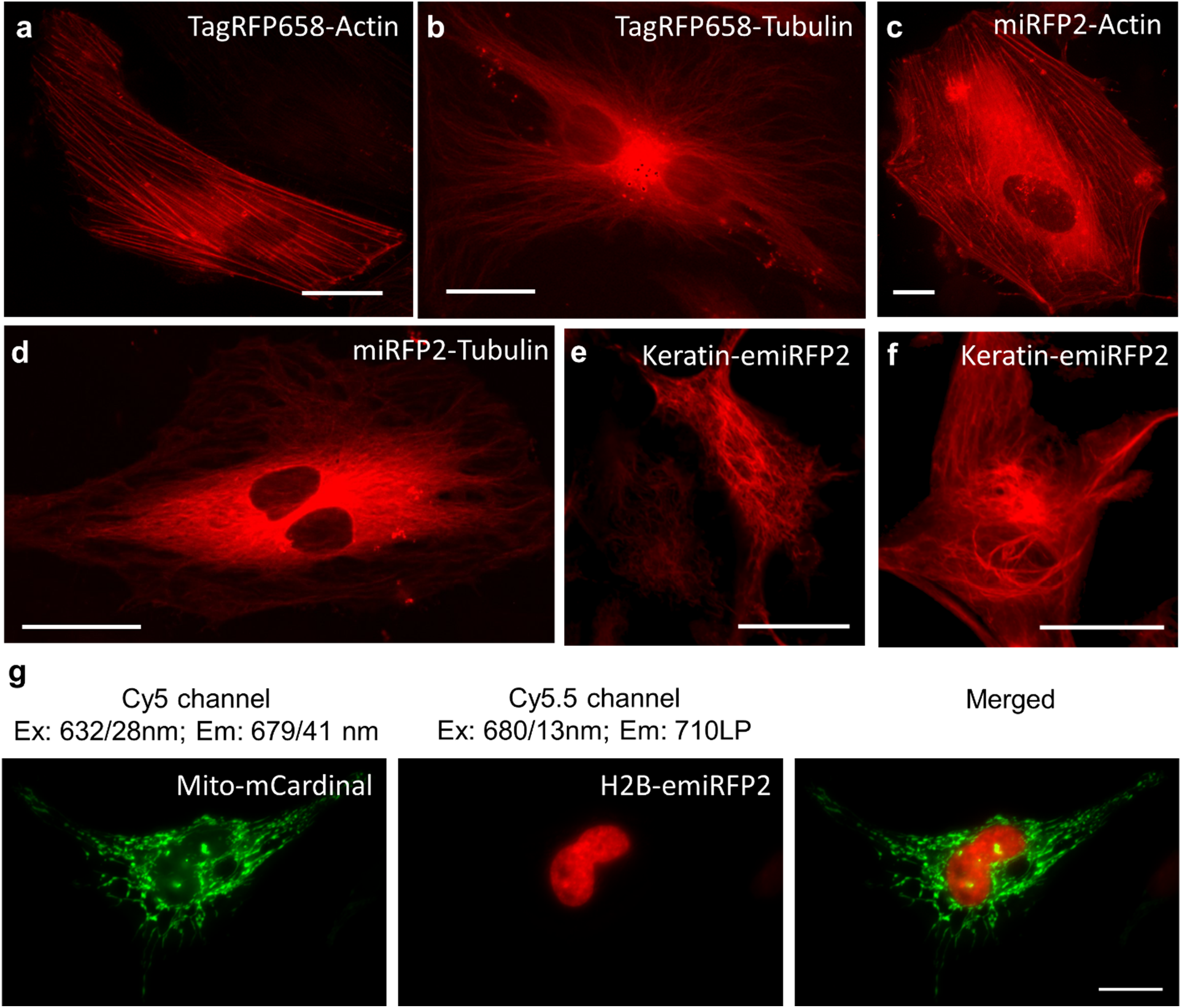
Fluorescence imaging of TagRFP658, miRFP2, and emiRFP2 fusions expressed in HeLa cells. (**a-f**) Representative fluorescence images of live HeLa cells transfected with (**a**) TagRFP658-α-Tubulin (n=11 cells from two independent transfections), (**b**) TagRFP658-β-Actin (n = 19 cells from two independent transfections). (**c**) miRFP2-α-Tubulin (n=15 cells from two independent transfections), (**d**) miRFP2-β-Actin (n=21 cells from two independent transfections), (e,f) Keratin-emiRFP2 (n=8 cells from two independent trasfections). Imaging conditions: (**a-d**) excitation 631/28 nm from an LED, emission 664LP under wide-field microscope. (**e,f**) excitation 640 nm from laser, emission 670-750 nm under confocal microscope. (**g**) Dual color NIR imaging of live HeLa cells co-expressing Mito-mCardinal and H2B-emiRFP2 (n=12 cells from two independent transfection)

## Discussion

Here we described a practical and simple approach for rapid optimization of FPs via directed molecular evolution in mammalian cells. The presented method does not require generation of stable cell lines or viral vector preparation and can be carried out using commonly available high-throughput cell sorting hardware such as FACS. One iteration of directed evolution accomplished with selecting individual variants can be performed in a course of ∼8 days, which is on average 5 to 8 times faster than alternative methods involving stable cell line generation^28,29,46^ or viral vector production^27,47,48^. The faster timeline compared to previously reported strategies was mainly achieved due to utilization of transient transfection of large gene libraries in combination with fast target gene recovery from small pools of collected cells (∼50-100 cells). Besides faster timeline, transient transfection provides additional advantages over traditional methods of gene library expression. It enables evolution of proteins for which it is impossible or very hard to establish stable cell lines, such as, for example, opsins^33^. In addition, it is more accessible than other methods as it only requires very standard expression vectors containing SV40*ori*, widely used HEK293T cells, and cheap transfection reagents. Besides clear advantages, there are also other experimental considerations that should be taken into account. Conditions of gene delivery are optimized that most of transfected cells (∼50%) receive single gene copy at total transfection efficiency of about ∼5%, which is similar to lentiviral transduction at a multiplicity of infection of ∼0.1 that is also used single gene copy delivery^27,49,50^. Therefore, in the case of large libraries it may significantly extend screening time as 95% of cells are negative. To reduce screening time using instrumental methods, negative cells can be removed using selective markers of choice, which can be included in the expression vector, although it will extend timeline as antibiotic selection usually takes 3-4 days or so. Furthermore, about 50% of transfected cells contain more than one gene^33^, which correspondingly requires screening of about twice more individual clones (Steps 7-9 of workflow in **Figure 1a**) than cells were collected during screening in order to ensure coverage of all diversity of recovered genes. However, it should be noted that lentiviral vectors provide similar efficiency of single gene copy delivery as gene delivery via both calcium phosphate transfection and viral transduction is a stochastic process with the same probability distribution. The developed method was tested only in HEK293T cells because this cell line provides sufficient expression level upon single copy plasmid transfection due to episomal replication of DNA facilitated by the SV40*ori* sequence in the expression vector. Besides HEK293T, there are several other commercially available cell lines expressing large T antigen required for episomal replication (available via ATCC) as well as self-replicating plasmids^51^, which can provide high enough expression level under required transfection conditions. However, applicability of other cell lines and expression vectors remains to be tested and verified. In this study we utilized FACS for high-throughput screening of mammalian cells, however any other method for fluorescence cells screening can be integrated into the described workflow. For example, SPOTlight approach^52^, microfluidc sorter^48^, photostick technique^53^, or robotic cell picker^33^ can be implemented following FACS enrichment of transfected cells, thus enabling phenotyping of single cells for a diversity of biochemical and photophysical properties of expressed mutated proteins, *e.g.* subcellular localization, pH stability, photostability, quantum yield etc.

The developed approach was validated by enhancing intracellular brightness of green and NIR FPs derived from evolutionary distinct naturally occurring chromoproteins. The starting template proteins, phiLOV2.1, TagRFP657, and miRFP, which were already highly optimized via directed molecular evolution either in *E.coli* or in mammalian cells, were significantly enhanced just in two iterations of directed evolution. We demonstrated that oligomerization or alternation of fluorescence spectra cannot be accounted for increase in intracellular brightness. In the case of TagRFP657 the enhancement of intracellular brightness can be most likely explained by the improvements in the chromophore maturation as TagRFP658 is characterized by higher extinction coefficient compared to its precursor. However, for phiLOV2.1 and miRFP the increase in intracellular brightness does not correspond to changes in molecular brightness. In contrast to GFP-like FPs, like TagRFP657, phiLOV2.1 and miRFP incorporate exogenous chromophores available in the cytoplasm of mammalian cells. Therefore, their intracellular brightness, besides of course intrinsic photophysical properties, will depend on efficiency of chromophore incorporation and chromophore availability. We revealed that miRFP2 has higher affinity towards BV than miRFP (**Supplementary Figure 8**), however it is hard to precisely quantify for which fraction of intracellular brightness enhancement the increase in affinity is responsible. The exact molecular basis of intracellular brightness increase observed for miRFP2 and phiLOV3 will require further investigations, which are beyond the scope of the present study. Overall, this study provided additional evidence, aligning with many previous reports, that relative molecular brightness of FPs expressed in *E.coli* does not always match intracellular brightness and the observed difference is particularly pronounced for the proteins that require exogenous chromophores. The presented results also justify the need for optimization of FPs in mammalian cells for higher performance *in vivo*.

To date, only four FPs were reported to be developed via directed molecular evolution in mammalian cells. Three of them are GFP-like FPs, mPlum^46^, Kriek^27^, and mCrispRed^28^, and one is BphP-based NIR FP miRFP^33^. A major rationale for evolving FPs in mammalian cells is to optimize their performance for in vivo applications, however, only mPlum and miRFP were tested in vivo. Crucially, side-by-side comparison of mPlum with other GFP-like FPs did not reveal any advantages of mPlum for in vivo imaging^54,55^, while miRFP was not compared to other BphP-based FPs either in cultured cells or in vivo. Therefore, it is important to demonstrate that the developed directed evolution approach capable of yielding practical FPs that outperform other FPs developed in *E.coli*. First, assessment of the developed FPs in HEK cells showed miRFP2 has comparable intracellular brightness to that of the best performing BphP-based NIR FPs reported to date, such iRFP670, miRFP680, and iRFP682. However, it should be noted that iRFP670, miRFP680, and iRFP682 although evolved in *E.coli* were actually selected in HeLa cells^37,56^. In turn rationally enhanced version of miRFP2 significantly outperformed other BphP-based NIR FPs in HEK cells and can be considered as the brightest BphP-based NIR FP in HEK cells (**Table 2**). Thus, the presented method enabled development of one of the best performing NIR FPs in HEK cells in the class just in 5 rounds of directed evolution (three rounds to select miRFP and two rounds to select miRPF2). However, further assessment of miRPF2 in vivo in model organisms reveal its inconsistent performance across various preparations. For example, miRFP2 co-expressed with HO1 was brighter mCardinal in fruit fly larvae, while dimmer in adult flies (**Figure 7h,i**). Fluorescence of miRFP2 in C.elegans can be enable only by HO1 co-expression. The intracellular brightness of emiRFP2 were 1.5-fold dimmer than mCardinal in cultured neurons, but about 4.8-fold dimmer in brain tissue (**Figure 6b** and **Figure 7b**). Furthermore, emiRFP2 was brighter than TagRFP658 in cultured neurons, while dimmer in zebrafish. We also observed significantly greater variation of Cy5-to-green fluorescence ratio for miRFP2 expressed in neurons in vivo in mice in comparison to cultured neurons, that is most likely due to higher variability of BV concentration in vivo. We suggest that high variability of miRFP2 performance in vivo in model organisms is determined by heme metabolism, which is required for BV production. For example, in wild type *C. elegans* heme utilization is not going via the BV intermediate as in vertebrates^57,58^, and thus expression of miRFP2 itself did not result in NIR fluorescence. However, we demonstrated that co-expression of HO1 is sufficient to enable BV production in *C.elegans*. Similarly, HO1 co-expression in Drosophila boosted fluorescence intensity of BphP-based NIR FPs expressed in neurons. Overall, these results indicated that high intracellular brightness of BphP-based NIR FPs in cultured cells may not be retained in vivo in model organisms, which complecats selection of the right NIR FP for in vivo applications and thus may require testing several candidates in real settings.

## Methods

### Molecular cloning and mutagenesis

The mammalian codon-optimized genes of phiLOV2.1 and UnaG were synthesized *de novo* by GenScript based on the amino acid sequences reported in the original publications^4,31^. The TagRFP657 and miRFP genes were acquired from Addgene (TagRFP657 plasmid#31959; miRFP plasmid#108409). Synthetic DNA oligonucleotides used for cloning were purchased from Integrated DNA Technologies. PrimeStar Max master mix (Clontech) was used for high-fidelity PCR amplifications. Restriction endonucleases were purchased from New England BioLabs and used according to the manufacturer’s protocols. Ligations were performed using T4 DNA ligase (Fermentas) or InFusion HD kits (Clontech). Small-scale isolation of plasmid DNA was performed with Mini-Prep kits (Qiagen); large-scale DNA plasmid purification was done with GenElute HP Endotoxin-Free Plasmid Maxiprep Kits (Sigma-Aldrich). Random libraries of mutants were prepared using Mutazyme II DNA polymerase (Agilent) under high mutation rate conditions (9-16 mutations per kilobase pair) and subcloned into the pN1 vector (Clontech). Obtained gene libraries in expression vectors were electroporated into NEB10-β *E.coli* host cells (New England BioLabs). Serial dilutions (10^−4^ and 10^−5^) of the electroporated cells were plated on LB/agar medium supplemented with 100 mg ml^−1^ kanamycin to estimate electroporation and cloning efficiency. For each library 20 randomly selected clones were sequenced to confirm ligation efficiency, *i.e.* fraction of the clones containing target genes, and mutation rate. The remainder of the cells was grown overnight in LB medium supplemented with 100 mg·ml^−1^ of kanamycin for subsequent plasmid DNA isolation. Library transfection into HEK293FT cells (Invitrogen) was performed as described previously. Briefly, HEK293FT cells were maintained between 10% and 70% confluence at 37 °C with 5% CO2 in DMEM medium (Cellgro) supplemented with 10% heat-inactivated FBS (Corning), 1% penicillin/streptomycin (Cellgro), and 1% sodium pyruvate (BioWhittaker). Transfection of HEK293FT cells with gene libraries was performed using a commercially available calcium phosphate (CaPhos) transfection kit (LifeTechnologies) according to the modified protocol using the pUC19 plasmid as “dummy” DNA in weight ratio library DNA: pUC19 1:100. To sort the gene library-transfected HEK293FT cells using flow cytometry, cells were harvested from a culture dish ∼ 48 h after gene library transfection by applying trypsin for 5–10 min (Cellgro) and then washed twice by centrifuging the cell suspension for 5 min at 500 rpm and re-suspending cells in PBS (Cellgro). The washed cells were then re-suspended in PBS supplemented with 4% FBS (Corning) and 10 mM EDTA at a density of 1–2·10^6^ cells/ml and filtered through a 30-µm filter (Falcon) to prevent clogging on the FACS machine. The filtered cells were sorted by FACSAria (BD Biosciences) running BD FACS Diva 8.0 software and equipped with standard 488- and 640-nm solid-state lasers. Debris, dead cells, and cell aggregates were gated out using forward and side scatter before desired fluorescence signals were detected. For each library, several hundred cells exhibiting highest fluorescent intensity in the corresponding channel were collected and subjected to whole genome amplification using a commercially available whole genome amplification kit (WGA, New England BioLabs) followed by PCR amplification. Amplicons with a size corresponding to that of the target gene were purified by agarose gel electrophoresis and cloned into the pN1 vector. Obtained colonies were individually picked to confirm the correct insert using PCR with a pair of primers that anneal to CMV promoter and the target genes. Colonies with correct insert were cultured in LB with 100 mg·ml^−1^ of kanamycin for plasmid purification. Purified plasmids were individually transfected into HEK293FT cells using TransIT-X2 reagent (Mirus). In 24 h post transfection, HEK cells were imaged using a wide-field Nikon Eclipse Ti equipped with 10x NA 0.45 and 20x NA 0.75 objective lenses, a SPECTRA-X light engine (Lumencor) with 475/28nm, and 631/28nm exciters (Semrock), a 5.5 Zyla camera (Andor), and automated stage (Ludl), controlled by NIS-Elements AR software Nikon to assess fluorescence and photostability.

The genes of miRFP720 and miRFP670nano were *de novo* synthesized by GenScript using sequences reported in the original publications^19,40^. The genes of mCardinal, mMaroon, and miRFP703 were acquired from Addgene as plasmids #54590, #54554, and #79988, respectively. To construct α-tubulin and β-actin fusions, the phiLOV3, TagRFP658, and miRFP2 genes were PCR amplified and swapped with mClover2 in pmClover2-tubulin-C-18 plasmid (Addgene #56376) and with TagRFP675 in pTagRFP675-actin-C1 (Addgene #44277) using InFusion cloning (Clontech). To construct Keratin fusions, emiRFP2 were PCR amplified and swapped with miRFP670nano in pKeratin-miRFP670nano plasmid (Addgene #127437) using Fusion cloning (Takara). To express TagRFP658 and mCardinal in neurons under CaMKII promoter the corresponding genes were PCR amplified and swapped with the Arch-GFP gene in FCK-Arch-GFP (Addgene #22217) using InFusion cloning. The pAAV-CAG-miRFP720-P2A-GFP and pAAV-CAG-miRFP2-P2A plasmids were constructed by cloning miRFP720-P2A and miRFP2-P2A into pAAV-CAG-GFP (Addgene #37825) in frame with GFP.

For expression in zebrafish larvae, the genes of mCardinal, TagRFP678, miRFP, and miRFP2 were codon-optimized using the online resource at http://www.bioinformatics.org/, synthesized *de novo* by GenScript. The zebrafish codon optimized emiRFP2 gene was cloned by substituting the nucleotides encoding *Rp*BphP1-based N-terminus of zebrafish codon optimized miRFP2 (aa 2-19) with those encoding 13 amino acid preceding the chromophore-binding Cys in *Rp*BphP2 as previously reported^20^.

For co-expression of TagRFP658, miRFP2, and emiRFP2 with mClover3 in NIH3T3 mouse embryonic fibroblasts (DSMZ) and PAC2 zebrafish embryonic fibroblasts, the zebrafish codon-optimized genes of the corresponding NIR FPs and mClover3 were cloned into an expression vector carrying the bidirectional promoter (p-EF1a:2xCMV:EF1a) consisting of two CpG free mCMV enhancers (Invivogen) each followed by CpG free hEF1alpha promoter (Invivogen) and SV40 polyA sequences. The generated expression cassettes were flanked by Tol2 transposon arms. The p-EF1α:2xCMV:EF1α bidirectional vector allows for expression of two reporter genes with comparable expression levels. For pan-neuronal expression in zebrafish, the zebrafish codon optimized gene of TagRFP658 was cloned into pTol2-UAS-zArchon1-KGC-GFP-ER2 plasmid (Addgene #108427) described before^33^ by swapping zArchon1-KGC-GFP-ER2 using SpeI and AscI sites.

For transient expression in cerebellar Purkinje neurons in zebrafish, EF1a:2xCMV:EF1a promoter in the above mentioned bidirectional vector carrying mClover3 and H2B-NIR FP, either of mCardinal, TagRFP658, or emiRFP2 was replaced by Purkinje neuron-specific bidirectional promoter (E1b:4xcpce:E1b) enabling co-expression of the corresponding NIR FPs with mClover3 predominantly in zebrafish cerebellar Purkinje cells (PCs) while also inducing slight expression in tectal neurons of larval zebrafish^45^.

For generation of transgenic flies, the mCardinal, mMarron, iRFP-VC, iRFP-VC-T2A-HO1, miRFP2, and miRFP2-T2A-HO1 genes were codon-optimized for expression in *Drosophila melanogaster* using the online resource at http://www.bioinformatics.org/, synthesized de novo and cloned into the 20XUAS-IVS-Jaws-mVenus_tr plasmid (Addgene #111553) by swapping the Jaws-mVenus gene.

For expression in *C.elegans*, the genes of miRFP2, miRP2-T2A-HO1, and jGCaMP7s were codon-optimized using SnapGene codon-optimization tool, synthesized *de novo* by GenScript and cloned into an expression vector under the tag-168 promoter the drives pan-neuronal expression.

### Protein purification and *in vitro* characterization

For protein purification, the phiLOV2.1, phiLOV3, TagRFP657, and TagRFP658 genes were cloned into pBAD-HisD vector and transformed into TOP10 cells. To express the protein, bacterial cells were grown in RM medium supplemented with ampicillin and 0.002% arabinose for 15–18 h at 37°C followed by 24 h at 18°C. Proteins were purified using TALON Metal Affinity Resin (Clontech) and dialyzed overnight against PBS buffer, pH 7.4. To express miRFP720 and miRFP2 the genes were cloned in pBAD-HisB plasmid and transformed into BW25113 bacterial cells, containing pWA23h plasmid encoding HO1 under rhamnose promoter. The colonies were grown in LB medium supplemented with ampicillin, kanamycin, 0.002% arabinose, and 0.02% rhamnose for 20 h at 37°C. For protein purification Ni-NTA agarose (Qiagen) was used. Protein was eluted with PBS containing 100 mM EDTA, followed by dialysis overnight against PBS buffer, pH 7.4.

Spectral properties of the proteins were measured in PBS at pH 7.4. The fluorescence and absorption spectra of phiLOV2.1, phiLOV3, TagRFP657, and TagRFP658 were measured using a Fluorolog3 spectrofluorometer (Jobin Yvon) and a Lambda 35 UV/Vis spectrometer (PerkinElmer), respectively. The fluorescence and absorption spectra of miRFP720 and miRFP2 were measured with a CM2203 spectrofluorometer (Solar, Belarus) and NanoDrop 2000c spectrophotometer (Thermo Scientific, USA), respectively. The extinction coefficient of phiLOV2.1 and phiLOV3 was determined as a ratio between the absorbance value of the peak at 374 nm, which correspond to FMN absorbance with extinction coefficient of 12,500 M^−1^·cm^−1^ (*ref.*^59^), and the value of the peak at major band peaked at ∼476 nm. For determination of the quantum yield, integrated fluorescence of phiLOV3 was compared with that of equally absorbing phiLOV2.1, characterized by quantum yield value of 0.2 (*ref.*^31^). To determine extinction coefficients of TagRFP657 and TagRFP658, we relied on measuring the mature chromophore concentrations, using alkali-denatured proteins as previously described^32^. For determination of the quantum yield, integrated fluorescence of TagRFP658 was compared with that of equally absorbing TagRFP657, characterized by quantum yield value of 0.1 (*ref.*^32^). The extinction coefficient of miRFP720 and miRFP2 was determined as a ratio between the absorbance value of the peak at Q-band and the value of the peak at Soret band, characterized by extinction coefficient of 39,900 M^−1^ cm^−1^ (*ref.*^60^). For quantum yield determination, the integrated fluorescence values of purified miRFP2 were compared with equally absorbing miRFP720 (quantum yield 0.061). pH titrations for phiLOV2.1, phiLOV3, TagRFP657, and TagRFP658 were performed in a 96-well black clear bottom plate (CORNING) by 1:20 dilution with a series of commercially available pH buffers (HYDRION) using a SpectraMax-M5 plate reader (Molecular Devices) to read out fluorescence. To determine pH stability, miRFP2 and miRFP720 were diluted 1:100 into a series of home-made pH adjusted buffers with NaOH and HCl (30 mM citric acid, 30 mM borax, or 30 mM NaCl) with pH values ranging from 3 to 10 in 0.5 pH units interval in a 96-well black clear bottom plate (Thermo Scientific, USA) using a Modulus^TM^ II Microplate Reader (TurnerBiosystems, USA) to read out fluorescence using 625nm excitation and 660-720 nm emission filters. Size exclusion chromatography for phiLOV3 and TagRFP658 was performed by GenScript on a Superdex 200 10/300 GL column (GE Healthcare Life Sciences) using a gel filtration standard (#1511901; BIO-RAD). Size exclusion chromatography for miRFP720 and miRFP2 was performed with a Superdex^TM^ 200 10/300 GL column using GE AKTA Explorer (Amersham Pharmacia, UK) FPLC System.

Two-photon spectrum and cross sections of TagRFP658 were measured in PBS buffer at concentrations ∼1–5·10^−5^ M in 1 mm glass spectroscopy cuvettes (Starna cells) using an MOM two-photon fluorescent microscope (Sutter Instrument) coupled with an Insight DeepSee (Newport) femtosecond laser, as described before^33^. For the measurement of spectral shape, fluorescence was collected through a combination of 694SP (Semrock) and 630/60 (Chroma) filters for TagRFP658 and through HQ705/100 (Chroma) filter for Styryl 9-M dye (Aldrich) in chloroform used as a reference standard. The two-photon cross section was measured relatively to LDS 698 dye (Exciton) in chloroform54 at 1150 nm using a combination of HQ705/100 (Chroma) and 630/60 (Chroma) filters and relatively to Rhodamine 700 in ethanol^61^ at 1200 nm using 675/20 (Chroma) filter. The spectral shape was then scaled to these values. The differences between the peak absolute values obtained with two different standards were within 10%.

### Protein characterization in mammalian cells

All procedures involving animals at MIT and Westlake University were conducted in accordance with the US National Institutes of Health Guide for the Care and Use of Laboratory Animals and approved by the Massachusetts Institute of Technology or Westlake University Committee on Animal Care. HEK293FT (Invitrogen) and HeLa (ATCC CCL-2) cells were maintained between 10% and 70% confluence at 37 °C with 5% CO_2_ in DMEM medium (Cellgro) supplemented with 10% heat-inactivated FBS (Corning), 1% penicillin/streptomycin (Cellgro), and 1% sodium pyruvate (BioWhittaker). Cells were authenticated by the manufacturer and tested for mycoplasma contamination to their standard levels of stringency and were here used because they are common cell lines for testing new tools. Primary mouse hippocampal neuronal culture was prepared as described previously. HEK293FT and HeLa cells were transiently transfected using TransIT-X2 (Mirus Bio LLC) or Calcium phospate transfection kit (K278001, Invitrogen) according to the manufacturer’s protocol and imaged 24-48 h after transfection. Mouse primary hippocampal neurons were prepared from postnatal day 0 or 1 Swiss Webster (Taconic) mice (both male and female mice were used) and cultured as previously described^33^. Neuronal cultures were transfected using Calcium phosphate transfection kit (Life Sciences) according to the protocol described before^33^. Transduction of neuronal culture was done at 4 DIV using ∼10^9^ viral particles of rAAV8-hSyn-miRFP2 (Janelia Farm Viral Core, the rAAV genome titer was determined by dot blot) per well of standard 24-well plate (CORNING). Imaging of HEK293FT cells and neuronal cultures for **Figure 1, 2** and **4** was done using a Nikon Eclipse Ti inverted microscope equipped with a SPECTRA X light engine (Lumencor) with 475/28 nm and 631/28 nm exciters (Semrock), and a 5.5 Zyla camera (Andor), controlled by NIS-Elements AR software, and using 10 × NA 0.3 and 40 × NA 1.15 objective lenses. Imaging of HEK293FT cells for **Figure 4** was done under using a Nikon Eclipse Ti2-E inverted microscope equipped with a SPECTRA III light engine (Lumencor) with 475/28 nm and 635/22 nm exciters (Semrock), a 680 nm solid state laser (MRL-III-680-800mW, CNI Laser) and 680/13 nm exciter (Semrock), and a Orca Flash4.0v3 camera (Hamamatsu), controlled by NIS-Elements AR software, and using 20 × NA 0.75 objective lenses. The efficiency of fluorescence imaging for the selected NIF FPs were calculated using the Microscope online application at www.fpbase.org using fluorescence spectra of the corresponding FPs (miRFP680 spectrum was substitute with iRFP682 spectrum, and miRFP2 and emiRFP2 spectra were substituted with miRFP spectrum due to their similarity; miRFP680 and miRFP2 spectra were not available at www.fpbase.org at the moment of data analysis), SpectraIII excitation profile, transmission spectrum of the emission filter, and quantum efficiency profile of the Octa-FlashV3 sCMOS camera.

NIH3T3 cells were grown in DMEM (4.5µg/L glucose) supplemented with 10% FCS, and maintained in a 5% CO_2_ incubator at 37°C. The NIH3T3 stable cell line was generated by the transfection of pEF-TGN46-mScarlet-iresPuro, followed by isolation and expansion of a clone grown in DMEM containing Puromycin at the concentration of 2µg/ml. Zebrafish PAC2 fibroblast cells were cultured in L-15 medium supplemented with 15% FCS, and maintained in an incubator at 28°C and atmospheric CO_2_. Transfection of plasmids was performed onto cells grown on µ-Slide 4-well glass bottom dish (ibidi) using jetPRIME reagent (Polyplus) according to the manufacturer’s protocol. For PAC2 cells, transfection was carried out in L-15 medium supplemented with 5% FCS. 6 hours after transfection, an equal volume of 15% FCS containing medium was added to adjust its final concentration of FCS at 10%. Live NIH3T3 and PAC2 cells were imaged 48 h post transfection using a laser scanning confocal microscope (TCS SP8, Leica Microsystems, Germany) with 40x water (N.A. 1.1) objective. The BV solution (Sigma) at the final concentration 25 μM was added to PAC2 cells 48 hours after transfection, followed by recording of their images 3 hours after the BV administration. mClover3 was excited by an argon laser at 488 mn and detection range at 495-530 nm, whereas each of NIR RFs was excited by HeNe 633 laser and detection range at 660-785nm. For three color imaging, transfected NIH3T3 cells were fixed with 4%PFA 36 hours after transfection, followed by imaging using a confocal microscope. mScarlet fluorescence was excited by a DPSS 561 nm laser with the detection range at 570-600 nm, followed by simultaneous imaging of mClover3 and the NIR FPs as described above.

### In utero electroporation and acute slice preparation

Embryonic day (E) 15.5 timed-pregnant female Swiss Webster mice (Taconic) were used for in utero electroporation as described before^33^. The pAAV-CAG-emiRFP2-P2A-GFP or pAAV-CAG-mCardinal-P2A-GFP plasmid at ∼1µg/µl concentration were injected into the lateral ventricle of one cerebral hemisphere of an embryo. Acute brain slices were obtained from mice at P20–30 without regard for sex using standard techniques as described before^33^. The brain slices were imaged using a Nikon Eclipse Ti2-E inverted microscope equipped with a SPECTRA III light engine (Lumencor) with 475/28 nm and 635/22 nm exciters (Semrock), and a Orca Flash4.0v3 camera (Hamamatsu), controlled by NIS-Elements AR software, and using 10x NA 0.45 and 20x NA 0.75 objective lenses.

### Zebrafish larvae preparation and imaging

All experiments involving zebrafish at MIT and Technische Universität Braunschweig were conducted in accordance with protocols approved by Massachusetts Institute of Technology Committee on Animal Care following guidelines described in the US National Institutes of *Health Guide for the Care and Use of Laboratory Animals* or by German legislation following European Union guidelines (EU Directive 2010_63) according to location of the respective experiments. Zebrafish larvae expressing TagRFP658 were prepared and imaged as described previously^33^. Briefly, pTol2-UAS-zTagRFP658 plasmid was co-injected with Tol2 transposase mRNA into homozygous nacre embryos of the pan-neuronal expressing Gal4 line, tg(elavl3:GAL4-VP16)nns6 (*ref.*^62^). Injected larvae were screened for near-infrared fluorescence in the brain and spinal cord at 2–3 d post fertilization using the Nikon wide-field microscope described above and subsequently imaged at 3–4 dpf using Zeiss Lightsheet Z.1 microscope. Lightsheets were generated by two illumination objectives (10×, NA 0.2), and the fluorescence signal detected by a 20×water immersion objective (NA 1.0). The laser line used for excitation was 638nm. Optical filters used to separate and clean the fluorescence response included a Chroma T647lpxr as a dichroic, and a Chroma ET665lp. Tiled data sets were taken with the Zeiss ZEN Software, and subsequently merged and processed with FIJI, and Arivis Vision4D.

To co-express the mCardinal, TagRFP658, and emiRFP2 H2B fusions with mClover3, one cell stage embryos of the pigmentation-compromised zebrafish *brass* strain and *Tg(4xen.cpce-E1B:gap-mScarlet)* line were microinjected with the corresponding E1b:4xcpce:E1b expression vector described above together with *tol2* mRNA (1.5 nl of injection mix containing 25 ng/μl of both *tol2* and pTol2-plasmid). mClover3 positive larval fish at 4 days post fertilization were subjected to confocal microscopy analysis. The stable transgenic line, *Tg(4xen.cpce-E1B: gap:mScarlet)* was generated by the injection of *tol1* mRNA together with a Tol1-reporter plasmid in which membrane-targeted mScarlet expression regulated by 4xcpce:E1b is restrictively induced in PCs, because ectopically expressed mScarlet in tectal neurons was eliminated by 4x miRNA181a target sequence inserted into the 3′UTR of the reporter gene^45^. For membrane targeting, mScarlet was fused N-terminally to the first 20 amino acids encoded by the zebrafish *gap43* gene. Fish larvae exhibiting no fluorescence in the corresponding channels were excluded from further imaging. Fluorescence imaging was performed using a laser scanning confocal microscope (TCS SP8, Leica Microsystems, Germany) with 40x water (N.A. 1.1) objectives. mClover3 (excited by an argon laser at 488 nm, detection range at 495-530 nm) and each of NIR FPs (excited by HeNe 633 nm laser, detection range at 660-785 nm) expressed in PCs were imaged simultaneously. For three-color imaging, first a DPSS 561 nm laser was used to excite mScarlet fluorescence, which was detected with the range at 570-600 nm, followed by simultaneous imaging of mClover3 and the NIR FPs as described above. Reconstructions and projections from z-stacks of images were generated with the 3D-projection program included in the LAS X software (Leica Microsystems, Germany). Acquired images were processed with FIJI to measure the fluorescent intensity ratio of each of NIR FPs and mClover3 in each PC. *In vivo* photostability of TagRFP658, emiRFP2, and mCardinal was assessed using PCs continuously exposed to HeNe633 laser set at 70% of the laser power in the software setting. The region of interest (116.25µm x 116.25µm) was drawn encompassing several PCs labeled with nuclear localized TagRFP658, or mCardinal, and single plan image (optical section: 3.56µm) was taken every 0.648 second without interval for 5 minutes. Acquired images were processed with FIJI to measure the fluorescent intensity in each PC in each time point.

### Preparation and imaging in Drosophila

Transgenic fly lines with the following genotypes y1 w67c23; P{y[+t7.7] w[+mC]=UAS-mCardinal} attP40, y1 w67c23; P{y[+t7.7] w[+mC]=UAS-mMaroon} attP40, y1 w67c23; P{y[+t7.7] w[+mC]=UAS-iRFP-VC} attP40, y1 w67c23; P{y[+t7.7] w[+mC]=UAS-iRFP-VC-T2A-HO1} attP40, y1 w67c23; P{y[+t7.7] w[+mC]=UAS-miRFP2 } attP40, y1 w67c23; P{y[+t7.7] w[+mC]=UAS-miRFP2-T2A-HO1} attP40 were generated by Bestgene using the user provided 20XUAS-IVS plasmids described above. Flies were raised on standard cornmeal medium at room temperature. To drive pan-neuronal protein expression, generated transgenic adult male flies were mated with C155-GAL4 (pan-neural, a gift from Littleton lab at MIT) virgin females to generate heterozygous (C155-GAL4/+ or Y; Transgene/+).

Intact 3^rd^ instar larva and 2-4-day old adult flies were immobilized on the coverslip for further imaging. To drive protein expression in specific neurons, generated transgenic flies were crossed with R84C10-GAL4 to generate heterozygous progenies. Dissected brains from 5-6-day old adult females were used for further imaging. No larvae or adult flies, carrying the genes of target proteins, were excluded from the study. The wide-field Nikon microscope described above was used to acquire low magnification images and a Zeiss LSM 800 confocal microscope with Airyscan equipped with 631 nm laser for excitation for high magnification images.

### *C. elegans* preparation and imaging

Worms were cultured and maintained following standard protocols^63^. Transgenic worms with extrachromosomal array co-expressing miRFP2 or miRFP2-T2A-HO1 with jGCaMP7s pan-neuronally were generated by co-injection of the two plasmids tag-168::miRFP2 or tag-168::miRFP2-T2A-HO1 with tag-168::NLS-jGCaMP7s into N2 background worms as described before^64^. All plasmids were injected at 10 ng/µl. Worms exhibiting the highest green fluorescence in neurons were picked and mounted on 2% agarose pads on microscope slides, worms without green fluorescence were excluded from further imaging, immobilized with 5 mM tetramisole, covered by a coverslip, and imaged using a Zeiss LSM 800 confocal microscope with Airyscan equipped with 631 nm laser for excitation.

### Data analysis and statistics

Data were analyzed offline using NIS-Elements Advance Research software, Excel (Microsoft), OriginPro, ImageJ, the Microscope online application (https://www.fpbase.org/microscope), and Arivis Vision4D. Data collection for fluorescence spectra of NIR FPs were done using https://www.fpbase.org. All attempts at replication of the experiments were successful. We did not perform a power analysis, since our goal was to create a new technology; and as recommended by the NIH, “In experiments based on the success or failure of a desired goal, the number of animals required is difficult to estimate…” As noted in the aforementioned paper, “The number of animals required is usually estimated by experience instead of by any formal statistical calculation, although the procedures will be terminated [when the goal is achieved].” These numbers reflect our past experience in developing neurotechnologies. All attempts at replication of the experiments were successful. No randomization was used in the study. No blinding was used in the study.

## Data availability

All other data generated or analyzed during this study are available from the corresponding author on reasonable request. The gene sequences of the new proteins will be deposited to GenBank.

## Acknowledgments

We thank B. Trout and C. Sudrik from MIT for help with spectroscopic analysis of phiLOV3 and TagRFP658. We also thank Shoh Asano from MIT for the help with zebrafish imaging, Cuiyun Bu from TU Braunschweig for the help with zebrafish injections, Shaofeng Wu from Westlake University for the help with transgenic worms preparation, and Yi Sun and Baoyue Liu from Westlake University for the help with fruit flies imaging. We are grateful to S. Flavel from MIT for the *C.elegans* expression vector. This work was supported by start-up funding from the Foundation of Westlake University, National Natural Science Foundation of China grant 32050410298, 2020 BBRF Young Investigator Grant, and MRIC Funding 103536022023 to K.D.P., an internal grant of the National Research Center Kurchatov Institute №1056 from 02.07.2020 to F.V.S., RFBR grant №19-04-00395 to O.M.S., RSF grant №17-14-01256 to T.V.K., the German Research Foundation (DFG, K1949/7-2) Project 241961032 to R.W.K., by Lisa Yang, John Doerr, HHMI, and grants NIH R01DA029639, NIH R01MH12297101, NIH R01DA045549, NIH R01MH114031, NIH R43MH109332, NIH R01GM104948, and NSF Grant 1734870 grants to E.S.B, and NIH BRAIN program U24NS109107 grant for M.D.

## Author Contributions

K.D.P. and E.S.B. initiated the project and made high-level designs and plans. K.D.P. and E.E.J. developed the rapid directed molecular evolution approach. K.D.P. and E.E.J. developed phiLOV3 and TagRFP658. K.D.P. and S.B. developed miRFP2. K.N. designed and cloned emiRFP2. K.D.P. characterized phiLOV3 and TagRFP568 *in vitro* and together with E.E.J. and H.Z. performed their characterization in mammalian cells. M.D. measured two-photon absorption properties in solution. L.E. and K.N. perform protein characterization in NIH3T3 and PAC2 cells. E.E.J. and K.D.P. performed electrophysiology experiments in cultured neurons. O.M.S., D.A.K., T.V.R., and F.V.S. characterized miRFP2 *in vitro*. K.D.P., H.Z., and S.B. characterized miRFP2 and emiRFP2 in mammalian cells. Y.W. performed IUE, and K.D.P. and H.Z. performed acute brain slice imaging. K.D.P., E.E.J., L.E., K.N., and R.W.K. prepared zebrafish and performed their imaging. J.S. and H.T. prepared transgenic worms and X.X. imaged them. K.D.P., S.B., D.P., and W.W. prepared transgenic flies and performed their imaging. K.D.P. performed the statistical analysis. K.D.P. oversaw all aspects of the project, analyzed and interpreted the data, wrote the manuscript with contributions from all of the authors.

## Additional information

Supplementary information accompanies this paper at

