## Supplementary Information for "Rapid Directed Molecular Evolution of Fluorescent Proteins in Mammalian Cells"

**Supplementary Table 1.** Screening conditions for the selected FPs.

| **Protein template** | **Library size (independent clones)** | **Fraction of clones with target genes (%)** | **FACS screening conditions** | **Fluorescence imaging conditions** |
| --- | --- | --- | --- | --- |
| phiLOV2.1 | 7.8·10^5^ | 92 | Ex: 488 nm;  Em: 515/20BP | 10x 0.45NA;  Ex: 475/34BP;  Em: 527/50BP; |
|  | 2.3·10^6^ | 100 |  |  |
| UnaG | 2.2·10^6^ | 83 | Ex: 488 nm;  Em: 515/20BP | 10x 0.45NA;  Ex: 475/34BP;  Em: 527/50BP; |
|  | 4.0·10^6^ | 100 |  |  |
| TagRFP657 | 1.1·10^6^ | 75 | Ex: 640 nm;  Em: 670/30BP | 10x 0.45NA;  Ex: 628/31BP;  Em: 664LP |
|  | 1.01·10^7^ | 100 |  |  |
| miRFP | 2.01·10^7^ | 95 | Ex: 640 nm;  Em: 710/50BP | 10x 0.45NA;  Ex: 628/31BP;  Em: 664LP |
|  | 1.8·10^6^ | 60 |  |  |

Ex – excitation wavelength; Em – emission wavelength; BP – bandpass; LP – longpass.

**Supplementary Figure 1.** Screening of the mutants selected in the final round of directed molecular evolution of UnaG, phiLOV2.1, TagRFP657, and miRFP.

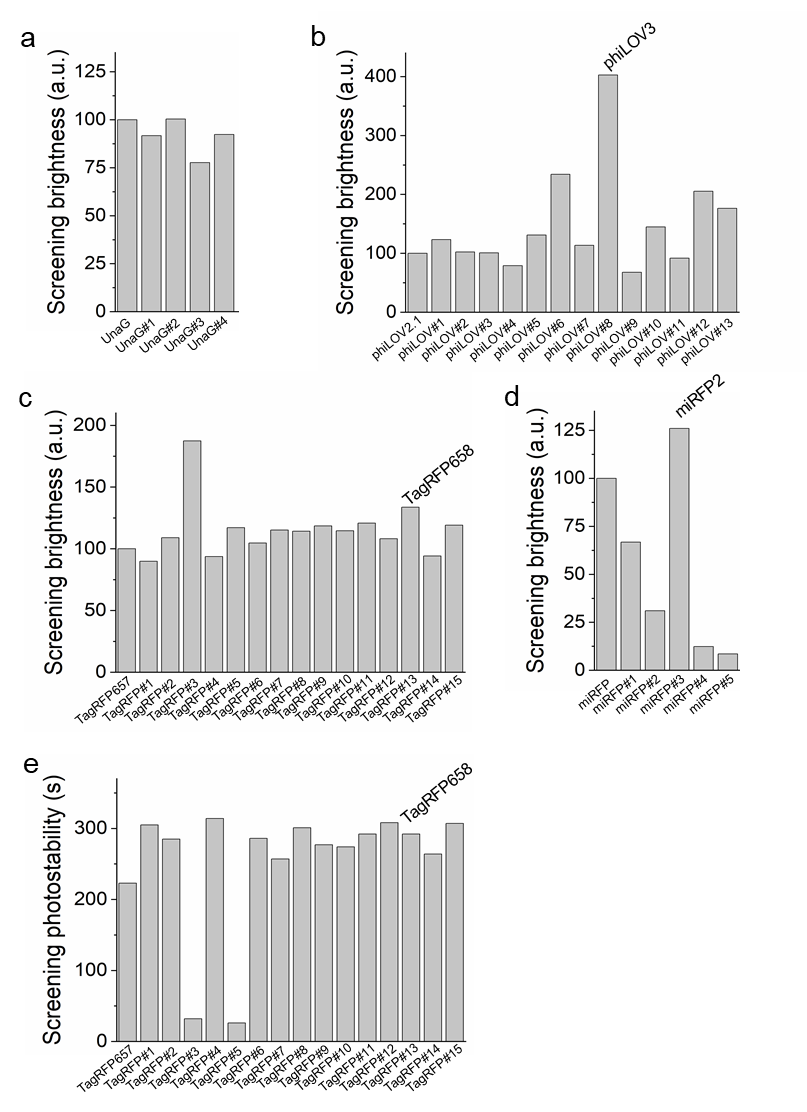

To assess fluorescence and photostability, the selected mutants were expressed in HEK cells under CMV promoter and imaged under wide-field microscope. (**a**) Screening fluorescence brightness of the top four mutants in comparison to their parental protein UnaG. (**b**, **c**, **d**) Screening fluorescence brightness of the selected (**b**) phiLOV2.1, (**c**) TagRFP657, and (**d**) miRFP mutants in comparison to the corresponding parental proteins. (**e**) Screening photostability of the selected TagRFP657 mutants in comparison to the parental protein TagRFP657.

**Supplementary Figure 2.** Alignment of amino acid sequences of UnaG and two of its selected mutants.

```
                 10        20        30        40        50        60
                 |         |         |         |         |         |
UnaG    MVEKFVGTWKIADSHNFGEYLKAIGAPKELSDGGDATTPTLYISQKDGDKMTVKIENGPP
UnaG#2  MVEKFVGTWKIADSHNFGEYLKAIGAPKELSGGGDATTPTLYISQKDGDKMTVKIENGPP
UnaG#4  MVEKFVGTWKIADSHNFGEYLKAIGAPKELSGGGDATTPTLYISQKDGDKMTVKVENGPP
```

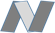

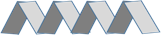

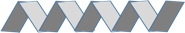

```
                 70        80        90        100       110       120
                 |         |         |         |         |         |
UnaG    TFLDTQVKFKLGEEFDEFPSDRRKGVKSVVNLVGEKLVYVQKWDGKETTYVREIKDGKLV
UnaG#2  TFLDTQVKFKLGEEFDEFPSDRRKGVKSVVNLVGEKLVYVQKWDGKETTYVREIKDGKLV
UnaG#4  TFLDTQVKFKLGEEFDEFPSDRRKGVKSVVNLVGEKLVYVQKWDGKETTYVREIKDGKLV
```

```
                 130
                 |
UnaG    VTLTMGDVVAVRSYRRATE
UnaG#2  VTLTMGDVVAVRSYRRATE
UnaG#4  VTLTMGDVVAVRSYRRATE
```

The chromophore-surrounding residues within 3.0 Å are highlighted in cyan (PDB: 4I3B). Mutant numbering corresponds to that in **Supplementary Figure 1a**. Introduced mutations are highlighted in red. The β-sheet-forming regions and α-helixes are shaded and denoted with arrows and ribbons, respectively.

**Supplementary Figure 3.** Alignment of amino acid sequences of phiLOV2.1 and phiLOV3.

```
                  10        20        30        40        50        60
                  |         |         |         |         |         |
phiLOV2.1 MEKSFVITDPRLPDYPIIFASDGFLELTEYSREEIMGRNARFLQGPETDQATVQKIRDAI
phiLOV3   MEKSFVITDPRLPDYPIIFASDGFLELTEYSREEIMGRNARFLQGPETDQATVQKIRDAI
```

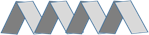

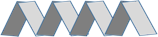

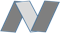

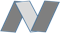

```
                  70        80        90        100       110
                  |         |         |         |         |
phiLOV2.1 RDQRETTVQLINYTKSGKKFWNLLHLQPVRDRKGGLQYFIGVQLVGSDHV
phiLOV3   RDRRETTVQLINYTKSGKKFWNLLHLQPVRDGKGGLQYFIGVQLVGSDHV
```

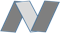

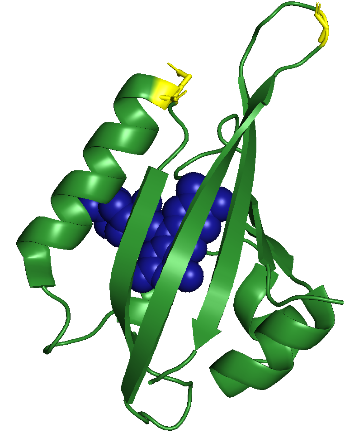

(*Top*) The residues surrounding the chromophore within 3.5 Å are highlighted in cyan (PDB: 4EEU). Mutations resulting in the conversion of parental phiLOV2.1 into the phiLOV3 variant are highlighted in red. The β-sheet-forming regions and α-helixes are shaded and denoted with arrows and ribbons, respectively. (*Bottom*) 3D protein structure of phiLOV2.1 (PDB 4EEU), chromophore shown in blue, amino acids mutated in phiLOV3 are shown in yellow.

**Supplementary Figure 4.** Alignment of amino acid sequences of TagRFP657 and TagRFP658.

```
                  10        20        30        40        50        60
                   |         |         |         |         |         |
TagRFP657 MSELITENMHMKLYMEGTVNNHHFKCTSEGEGKPYEGTQTQRIKVVEGGPLPFAFDILAT
TagRFP658 MSELITENMHMKLYMEGTVNNHHFKCTSEGEGKPYEGTQTQRIKVVEGGPLPFAFDILAT
```

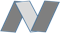

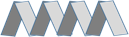

```
                  70        80        90       100       110       120
                   |         |         |         |         |         |
TagRFP657 SFMYGSHTFINHTQGIPDFWKQSFPEGFTWERVTTYEDGGVLTATQDTSLQDGCLIYNVK
TagRFP658 SFMYGSHTFIDHTQGIPDFWKQSFPEGFTWERVTTYEDGGVLTATQDTSLHDGCLIYNVK
```

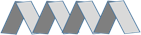

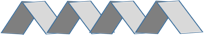

```
                 130       140       150       160       170       180
                   |         |         |         |         |         |
TagRFP657 IRGVNFPSNGPVMQKKTLGWEAHTEMLYPADGGLEGRTALALKLVGGGHLICNFKTTYRS
```

**TagRFP658 IRGVNFPSNGPVMQKKTLGWEAHTEMLYPTDGGLEGRTALALKLVGGGHLICNFKTTYRS**

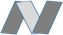

```
                 190       200       210       220       230
                   |         |         |         |         |
TagRFP657 KKPAKNLKMPGVYYVDYRLERIKEADKETYVEQHEVAVARYCDLPSKLGHKLN
TagRFP658 KKPAKNLKMPGVYYVDYRLERIEEADNETYVEQHEVAVARYCDLPTKLGHKLN
```

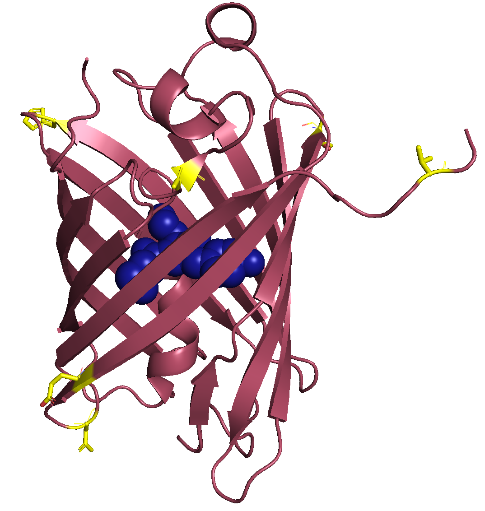

(*Top*) Internal amino acids are shaded, and the chromophore-forming residues are underlined. Mutations resulting in the conversion of TagRFP657 into TagRFP658 are highlighted in red. The β-sheet-forming regions and α-helixes are shaded and denoted with arrows and ribbons, respectively. (*Bottom*) 3D protein structure of TagRFP (PDB 3M22, precursor of TagRFP657), chromophore shown in blue, amino acids mutated in TagRFP658 are shown in yellow.

**Supplementary Figure 5.** Alignment of amino acid sequences of PAS-GAF domains of the *Rp*BPhP1 and its derivatives miRFP and miRFP2.

```
                  10        20        30        40       50        60
                  |         |         |         |         |         |
RpBPhP1  MVAGHASGSPAFGTADLSNCEREEIHLAGSIQPHGALLVVSEPDHRIIQASANAAEFLNL
miRFP    MVAGHASGSPDFGTADPSDCEREEIHLAGSIQPHGTLLVVSEPDHRIIQASANAAEFLNL
miRFP2   MVAGHASGSPDFGTASPSDCEREEIHLAGSIQPHGTLLVVSEPDHRIIQASANAAEFLNL
```

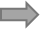

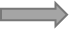

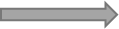

```
                 70        80        90        100       110       120
                  |         |         |         |         |         |
RpBPhP1  GSVLGVPLAEIDGDLLIKILPHLDPTAEGMPVAVRCRIGNPSTEYDGLMHRPPEGGLIIE
miRFP    GSVLGVPLAEIDGDLLIKILPHLDPTAEGMPVAVRCRIGNPSTEYDGLMHRPPEGGLIIE
miRFP2   GSVLGIPLAEIDGDLLIKILPHLDLTAEGMPVAVRCRIGNPSTEYDGLMHRPPEGGLIIE
```

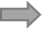

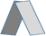

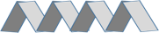

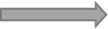

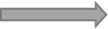

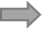

```
                 130       140       150       160       170       180
                  |         |         |         |         |         |
RpBPhP1  LERAGPPIDLSGTLAPALERIRTAGSLRALCDDTALLFQQCTGYDRVMVYRFDEQGHGEV
miRFP    LERAGPPIDLSGTLAPALERIRTAGSLRALCDDTALLFQQCTGYDRVMVYRFDEQGHGEV
miRFP2   LERAGPSIDLSGTLAPALERIRTAGSLRALCDDTVLLFQQCTGYDRVMVYRFDERGHGEV
```

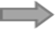

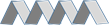

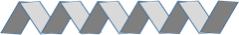

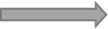

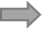

```
                 190       200       210       220       230       240
                  |         |         |         |         |         |
RpBPhP1  FSERHVPGLESYFGNRYPSSDIPQMARRLYERQRVRVLVDVSYQPVPLEPRLSPLTGRDL
miRFP    YSEIHVTGLESYFGNRYPSSLVPQMARRLYERQRVRVLVDVSYQPVPLEPRLSPLTGRDL
miRFP2   YSEIHVTGLESYFGNRYPSSLVPQMARRLYVRQRVRVLVDVTYQPVPLEPRLSPLTGRDL
```

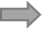

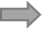

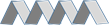

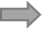

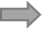

```
                 250       260       270       280       290       300
                  |         |         |         |         |         |
RpBPhP1  DMSGCFLRSMSPIHLQYLKNMGVRATLVVSLVVGGKLWGLVACHHYLPRFIHFELRAICE
miRFP    DMSGCFLRSMSPTHLQFLKNMGVRATLVVSLVVGGKLWGLVICHHYLPRFIHFELRAICE
miRFP2   DMSGCFLRSMSPTHLQFLKNMGVRATLVVSLVVGGKLWGLVICHHYLPRFIHFELRAICV
```

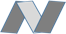

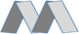

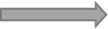

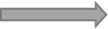

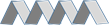

```
                 310
                  |
RpBPhP1  LLAEAIATRITAL
miRFP    LLAEAIATRITAL
miRFP2   LLAEAIATRITAL
```

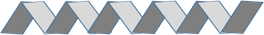

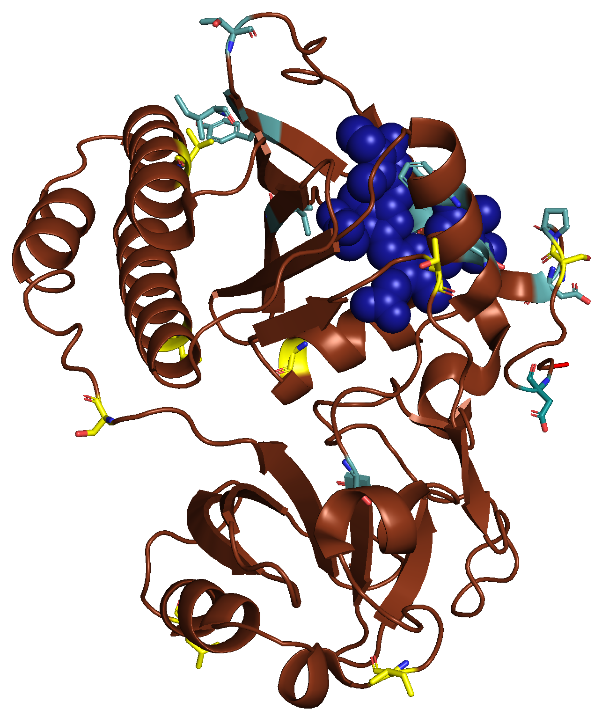

(*Top*) The residues surrounding the chromophore within 4.0 Å in *Rp*BPhP1 are highlighted in cyan (PDB: 4GW9). Mutations resulting in the conversion of parental *Rp*BphP1 into miRFP are highlighted in green. Mutations resulting in the conversion of miRFP into miRFP2 variant are highlighted in red. The β-sheet-forming regions and α-helixes are shaded and denoted with arrows and ribbons, respectively. (*Bottom*) 3D protein structure of RpBphP1 photosensory core domain (PDB 5OY5; precursor of miRFP), chromophore shown in blue, amino acids mutated in miRFP are shown in cyan, amino acids mutated in miRFP2 are shown in yellow.

**Supplementary Figure 6.** Two-photon cross-section spectrum of TagRFP658.

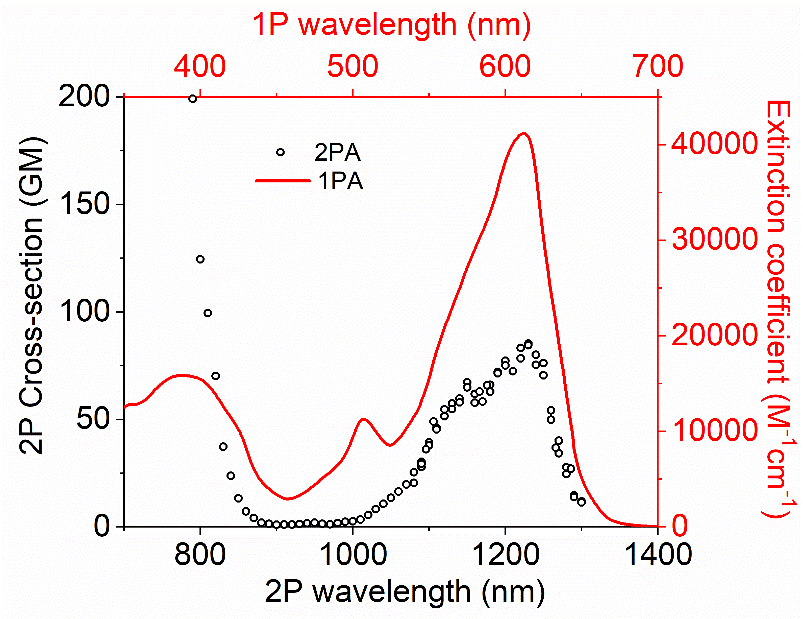

One-photon absorption (solid line) and two-photon cross-section (open circles) absorption spectra of TagRFP658. Two-photon cross-section is presented versus laser wavelength used for excitation. GM, Goeppert-Mayer units.

**Supplementary Figure 7.** Size exclusion chromatography calibration plots.

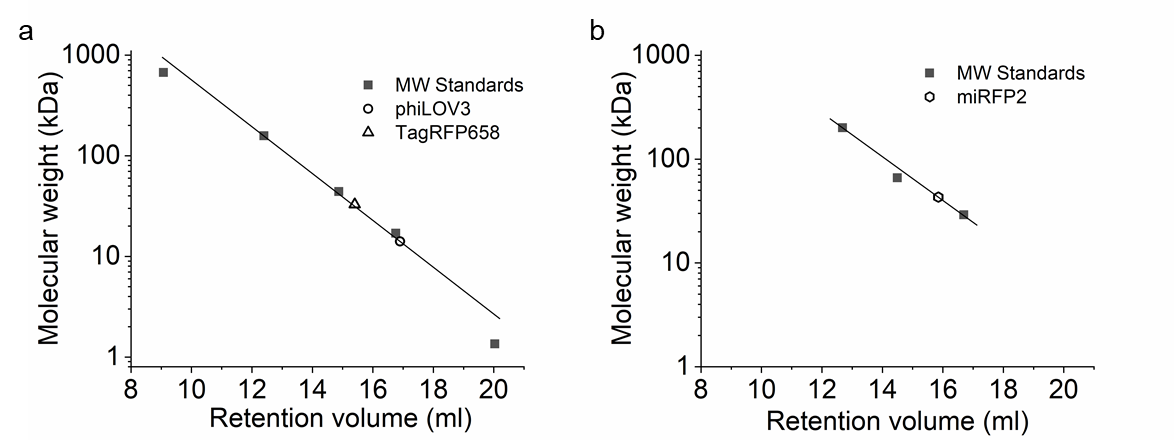

Size exclusion chromatography calibration plots showing the relative retention volumes of protein molecular weight standards and proteins of interest (see Figure 2 for size exclusion chromatography profiles). (**a**) Calibration plot showing the relative retention volumes of protein molecular weight standards (black squares; Gel Filtration Standard, Bio-Rad; n = 1 technical replicate) and phiLOV3 (circle) and TagRFP658 (triangle). (**b**) Calibration plot showing the relative retention volumes of protein molecular weight standards (black squares; n = 1 technical replicate) and miRFP2 (hexagon).

**Supplementary Figure 8.** Effect of exogenous biliverdin on NIR fluorescence of miRFP703, miRFP, and miRFP2.

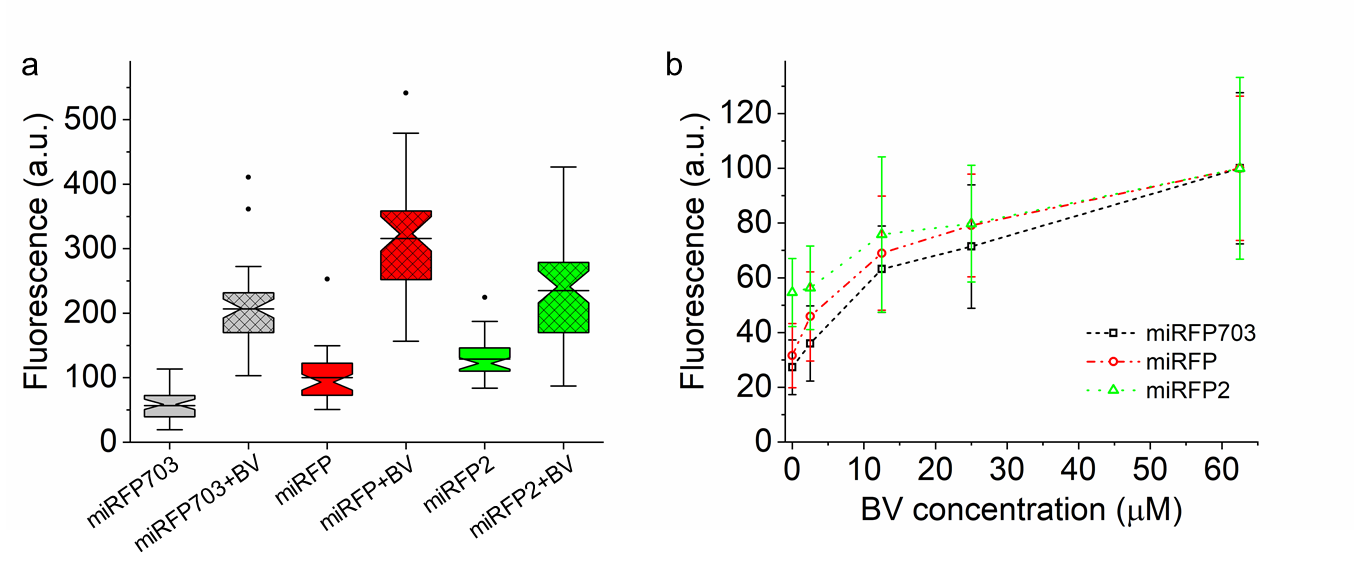

(**a**) Relative normalized fluorescence of HEK cells expressing miRFP703 (gray), miRFP (red), and miRFP2 (green) under CMV promoter with (cross-hatched boxes) and without (open boxes) addition of 62.5 µM for 3 h before imaging (n = 44, 45, 40, 40, 41, and 48 cells for miRFP703, miRFP703+BV, miRFP, miRFP+BV, miRFP2, and miRFP2+BV, respectively, from one independent transfection each). Imaging conditions: excitation  631/28 nm from an LED, emission 664LP. Box plots with notches are used (see caption for **Figure 1c** for description). (**b**) Exogeneous BV binding to miRFP703 (black dashed line), miRFP (red dashed line), and miRFP2 (green dashed line) proteins expressed in HEK cells. Cells were incubated with the respective concentration of BV for 3 h before imaging (n= ~40 cells for each protein under each condition from one independent transfection). Open symbols, mean; error bars, standard deviation. Imaging conditions same as in (**a**).

**Supplementary Figure 9.** Quantitative assessment of the selected near-infrared FPs in HEK cells.

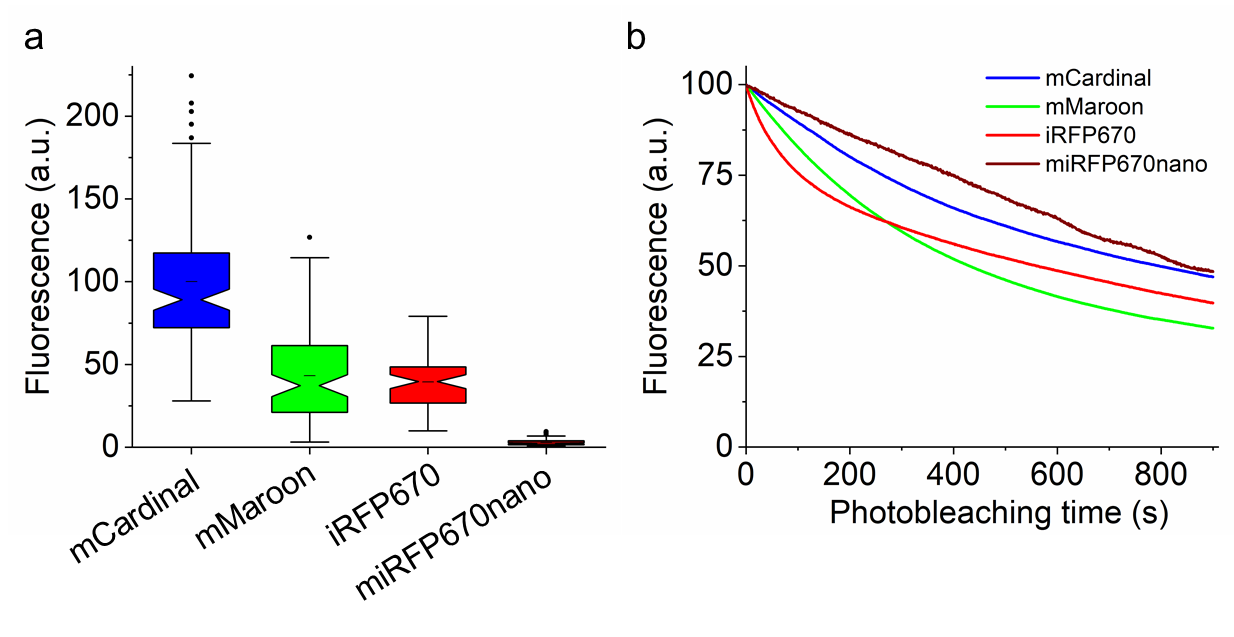

The selected FPs were expressed in HEK cells under CMV promoter and imaged 48 h post transfection. (**a**) Relative normalized fluorescence and (**b**) raw photobleaching curves for mCardinal (blue), mMaroon (green), iRFP670 (red), and miRFP670nano (deep red) expressed in HEK cells (for fluorescence n=127, 93, 65, and 119 cells, respectively, from one culture; for photobleaching n=19, 12, 38, and 10 cells, respectively, from one culture; imaging conditions: wide-field excitation 631/28 nm from LED at 25 mW/mm^2^, emission 664LP).

**Supplementary Figure 10.** Membrane properties of cultured primary mouse hippocampal neurons expressing mCardinal and TagRFP658.

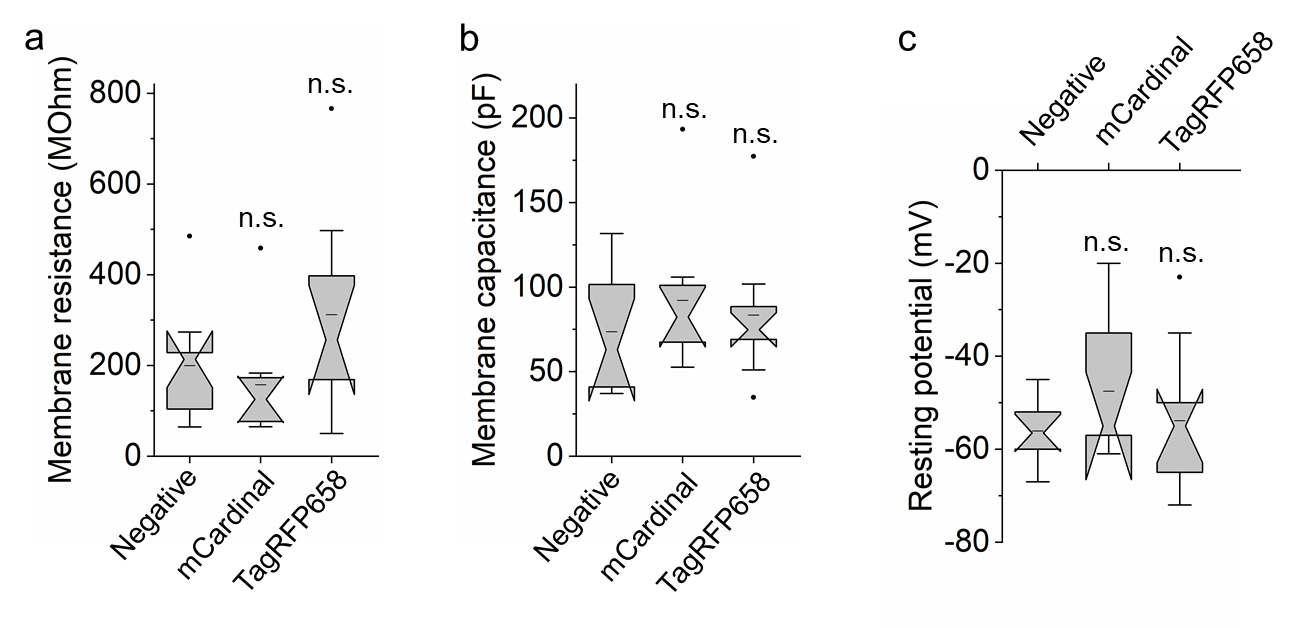

Cultured hippocampal neurons expressing mCardinal (n=9 cells from one culture) and TagRFP658 (n=9 cells from one culture) under CaMKII promoter were patched to compare membrane properties to non-transfected neurons (negative control, n=10 cells from two cultures). (**a**) Membrane resistance. P > 0.05, not significant (n.s.) Kruskal–Wallis one-way analysis of variance throughout all panels of this figure. (**b**) Membrane capacitance. (**c**) Resting potential. Throughout this figure, box plots with notches are used; narrow part of notch, median; top and bottom of the notch, 95% confidence interval for the median; top and bottom horizontal lines, 25% and 75% percentiles for the data; whiskers extend 1.5 times the interquartile range from the 25th and 75th percentiles; horizontal line, mean. For datasets with n < 10, open circles represent individual data points; data points which are less than the 25th percentile or greater than the 75th percentile by more than 1.5 times the interquartile range are also represented as open circles.

**Supplementary Figure 11.** Expression of miRFP2 in primary mouse hippocampal neurons.

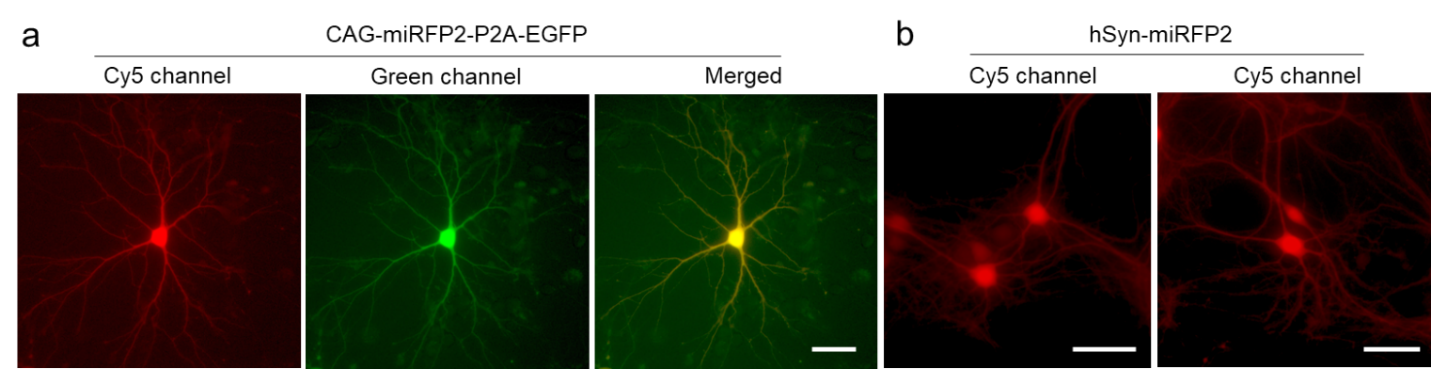

(**a**) Representative image of calcium-phosphate transfected mouse hippocampal neuron expressing miRFP2-P2A-EGFP under CAG promoter (n=40 neurons from one culture). Imaging conditions: NIR channel, excitation  631/28 nm from an LED, emission 664LP; green channel: excitation 475/34BP from an LED, emission 527/50 nm. (**b**) Representative images of rAAV-transduced mouse hippocampal neurons expressing miRFP2 under hSyn promoter (n=120 neurons from one culture). Imaging conditions: NIR channel, excitation  631/28 nm from an LED, emission 664LP.

**Supplementary Figure 12.** Intracellular brightness and photostability of TagRFP658, miRFP2 and emiRFP2 in live HEK cells in Cy5 and Cy5.5 channels.

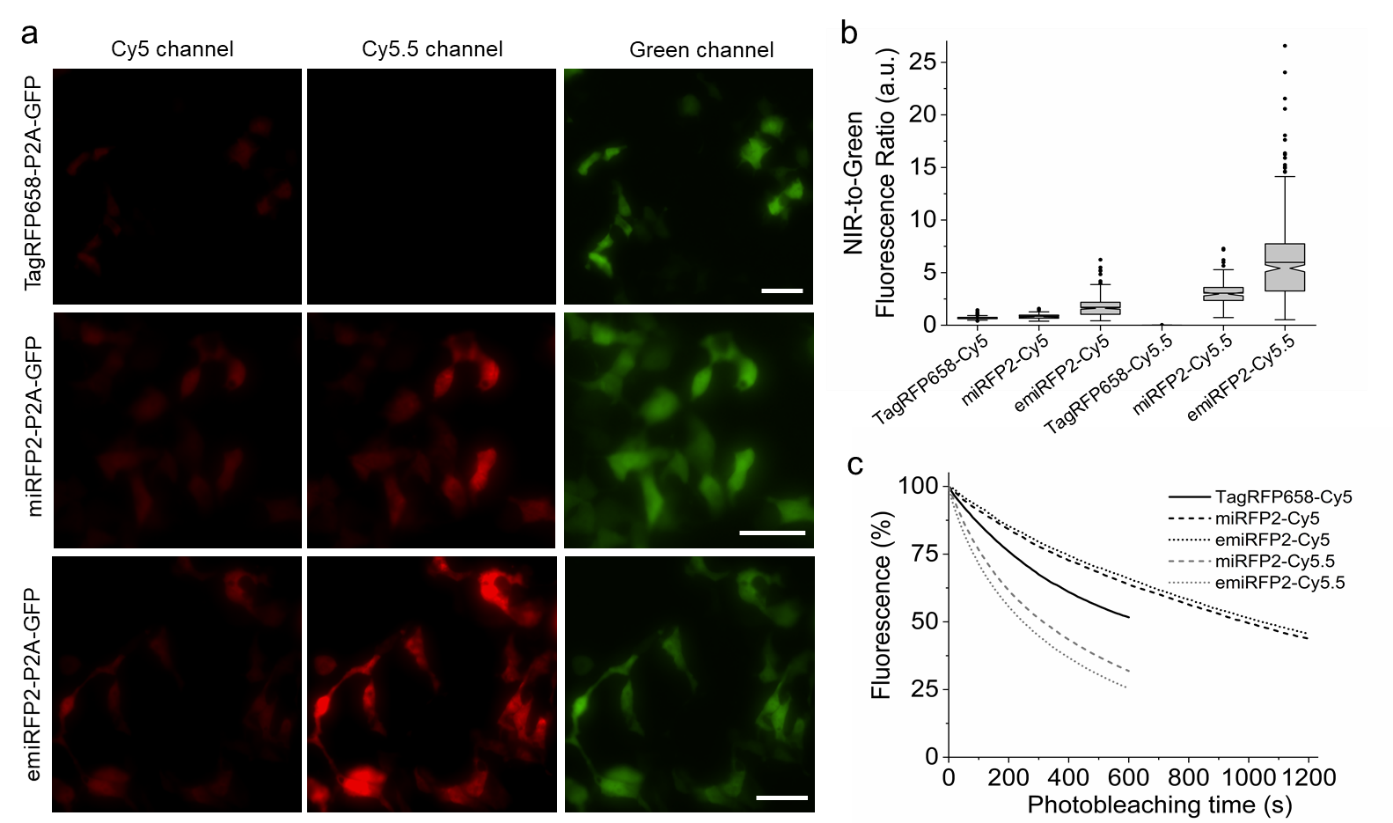

Intracellular brightness and photostability of TagRFP658, miRFP2, and emiRFP2 in live HEK cells. (**a**) Representative fluorescence images of cells transfected with pAAV-CAG-TagRFP658-P2A-EGFP (top), pAAV-miRFP2-P2A-EGFP (middle), and pAAV-emiRFP2-P2A-EGFP (bottom; n = 226, 131, and 475 cells from 2, 4, and 2 independent transfections, respectively). Imaging conditions: Cy5 channel: excitation 635/22 nm from 637 nm laser, emission 730/140 nm; Cy5.5 channel: excitation 680/13 nm from 680 nm laser, emission 710 LP; GFP channel: excitation 478/24 nm for an LED; emission 535/46 nm. Images in Cy5 and Cy5.5 were taken under matching excitation intensity (66 mW/mm2) and the same exposure time (100 ms). The dynamic range of fluorescence intensity in Cy5 and Cy5.5 channels are identical across all images. Scale bar, 50 µm. (b) NIR-to-green fluorescence ratio for TagRFP658, miRFP2, and emiRFP2 in live HEK cells shown in a (n = 226, 131, and 475 cells from 2, 4, and 2 independent transfections, respectively). Box plots with notches are used in this figure (see Fig. 1c for the description). (c) Intracellular photostability of TagRFP658, miRFP2, and emiRFP2 in Cy5 and Cy5.5 channels (n = 56, 57, and 37 cells for TagRFP658, miRFP2, and emiRFP2131, and 475 cells from 2 independent transfections each under Cy5 excitation; n = 44 and 78 from 2 and 3 independent transfection for miRFP2 and emiRFP2, respectively). Imaging conditions the same as in a.

**Supplementary Figure 13.** Imaging of miRFP2 in transgenic *C.elegans* worms and *Drosophila* *melanogaster.*

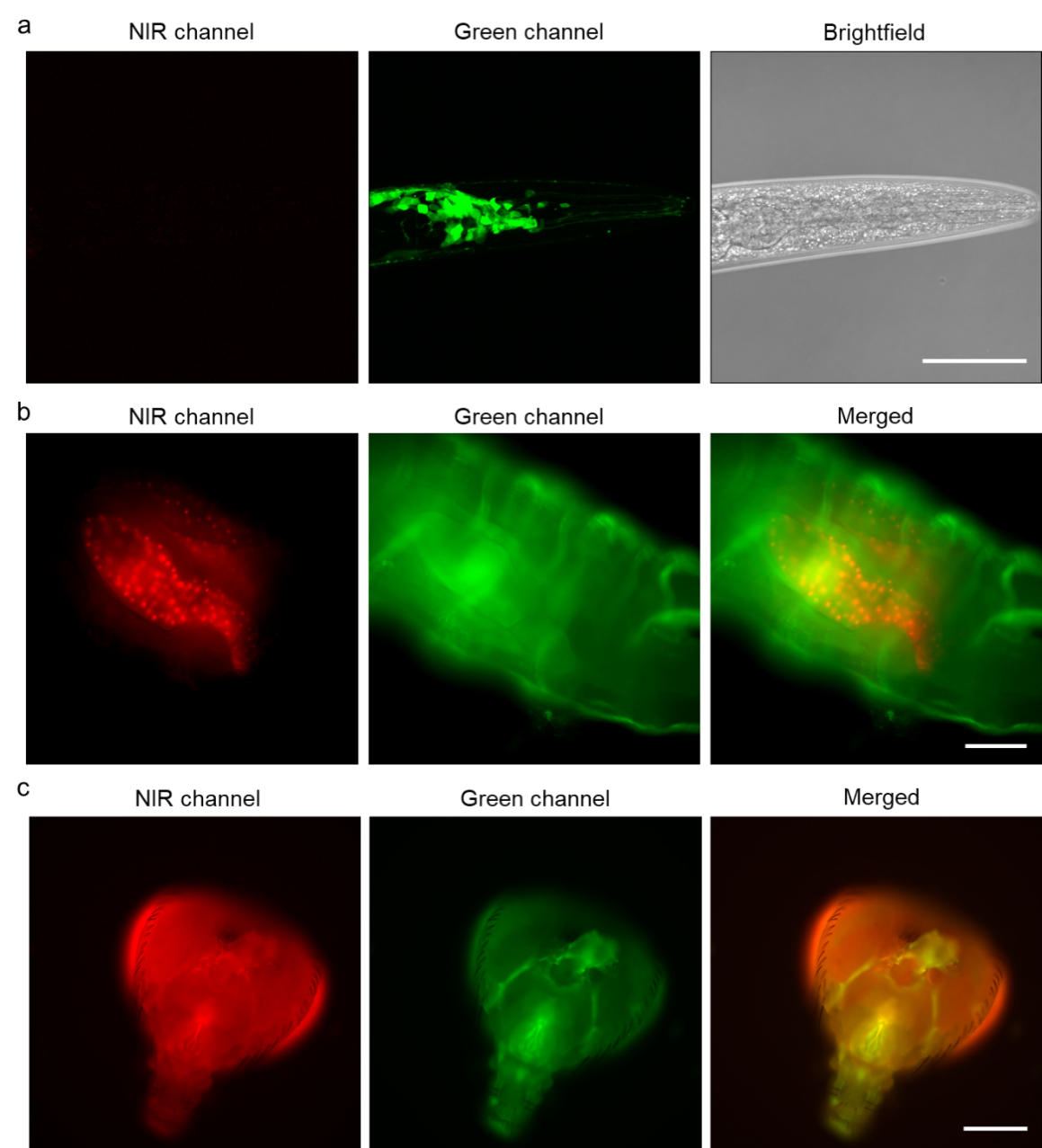

Live transgenic *C.elegans* worms and *Drosophila* larvae with pan-neuronal expression of corresponding codon-optimized genes of miRFP2 were imaged in NIR and green channels. (**a**) Representative fluorescence and brightfield images of the *C.elegans* head co-expressing codon-optimized genes of miRFP2 and jGCaMP7b in neurons (n=15 worms from two independent microinjections). Imaging conditions: NIR channel, excitation 631 nm from a laser, emission 645-700 nm; green channel, excitation 488 nm from a laser, emission 500-650 nm. Scale bar, 50 µm. (**b**) Representative fluorescence images of live intact 3^rd^ instar *Drosophila* larva expressing miRFP2 (n= 10 larvae from two transgenic lines). Imaging conditions: NIR channel, excitation 631/28 nm from LED, emission 665LP; green channel, exctitation  475/34 nm from LED, emission 527/50 nm (green fluorescence correspond to autofluorescence). Scale bar, 250 µm. (**c**) Representative fluorescence images of adult *Drosophila* fly head expressing miRFP2 (n= 11 flies from one transgenic line). Imaging conditions the same as in **b**. Scale bar, 250 µm.

**Supplementary Figure 14.** Influence of HO1 co-expression on fluorescence of miRFP2 in live HEK cells.

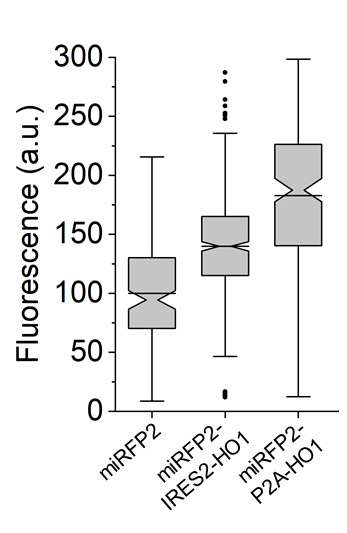

Relative fluorescence of live HEK cells expressing miRFP2, miRFP2-IRES2-HO1, and miRFP2-P2A-HO1 under CMV promoter (n= 142, 394, and 187 cells, respectively, from two independent transfections each).

**Supplementary Figure 15.** Triple color imaging of fixed NIH3T3 cells.

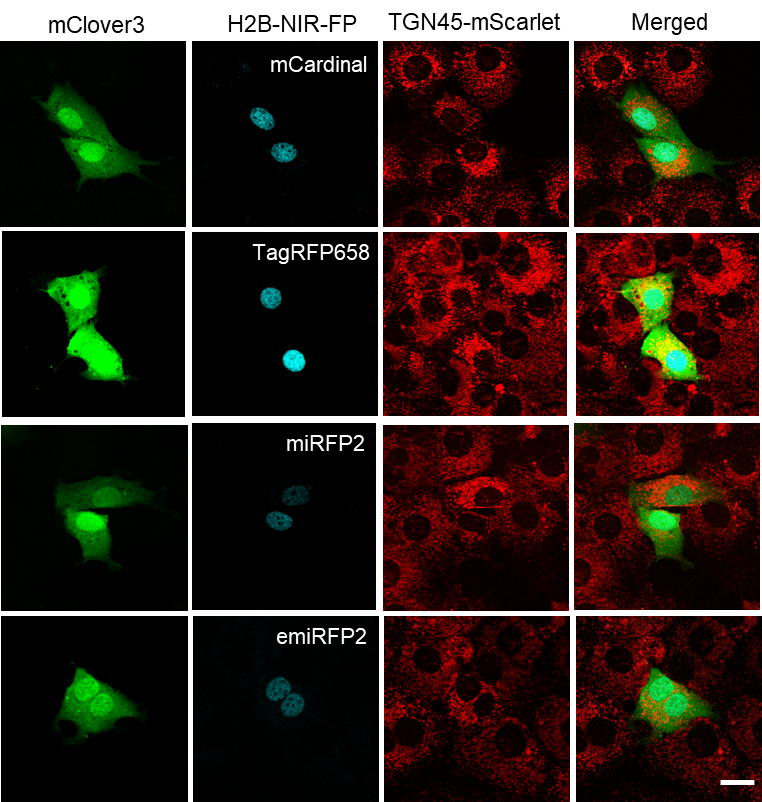

Each bidirectional expression construct carrying either of H2B fused with NIR-FP together with mClover3 was transfected into NIH3T3 cells stably expressing mScarlet N-terminally fused to TGN46 located in TGN and the vesicles cycling between TGN and the cell surface. Transfected cells were fixed by 4% PFA, followed by imaging using confocal microscope in green, red, and Cy5 channels. Imaging conditions: green channel, excitation 488 nm from an argon laser, emission 495-530 nm; Red channel, excitation 561 nm from a laser, emission 570-600 nm. Cy5 channel, excitation 633 nm from a laser, emission 660-785 nm. Scale bar, 20 µm.

**Supplementary Figure 16.** Dual-color *in vivo* imaging of wild type zebrafish larvae co-expressing NIR FPs with mClover3 in PCs.

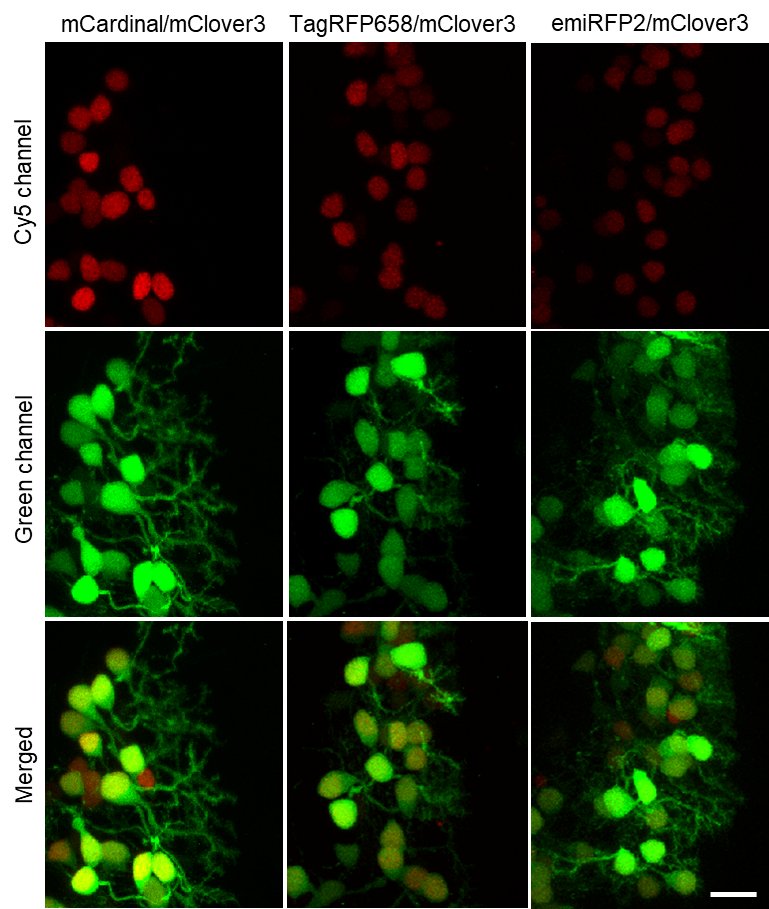

Representative confocal images of 4 dpf wild type zebrafish larvae transiently co-expressing H2B-NIR FPs with mClover3. Each subset of images (top to bottom) shows the co-expression of each H2B-NIR-FP fusion (left, mCardinal; middle, TagRFP658; right, emiRFP2) with cytoplasmic mClover3 (n=4 fish for each subset). Imaging conditions, Cy5 channel: excitation 633 nm laser, emission 722/63 nm; green channel: excitation 488 nm laser, emission 513/17 nm. Scale bar, 10 µm.
